# A metabotropic glutamate receptor agonist enhances visual signal fidelity in a mouse model of retinitis pigmentosa

**DOI:** 10.1101/2024.04.30.591881

**Authors:** Xiaoyi Li, Miloslav Sedlacek, Amurta Nath, Klaudia P. Szatko, William N. Grimes, Jeffrey S. Diamond

## Abstract

Many inherited retinal diseases target photoreceptors, which transduce light into a neural signal that is processed by the downstream visual system. As photoreceptors degenerate, physiological and morphological changes to retinal synapses and circuitry reduce sensitivity and increase noise, degrading visual signal fidelity. Here, we pharmacologically targeted the first synapse in the retina in an effort to reduce circuit noise without sacrificing visual sensitivity. We tested a strategy to partially replace the neurotransmitter lost when photoreceptors die with an agonist of receptors that ON bipolars cells use to detect glutamate released from photoreceptors. In *rd10* mice, which express a photoreceptor mutation that causes retinitis pigmentosa (RP), we found that a low dose of the mGluR6 agonist L-2-amino-4-phosphonobutyric acid (L-AP4) reduced pathological noise induced by photoreceptor degeneration. After making *in vivo* electroretinogram recordings in *rd10* mice to characterize the developmental time course of visual signal degeneration, we examined effects of L-AP4 on sensitivity and circuit noise by recording *in vitro* light-evoked responses from individual retinal ganglion cells (RGCs). L-AP4 decreased circuit noise evident in RGC recordings without significantly reducing response amplitudes, an effect that persisted over the entire time course of rod photoreceptor degeneration. Subsequent *in vitro* recordings from rod bipolar cells (RBCs) showed that RBCs are more depolarized in *rd10* retinas, likely contributing to downstream circuit noise and reduced synaptic gain, both of which appear to be ameliorated by hyperpolarizing RBCs with L-AP4. These beneficial effects may reduce pathological circuit remodeling and preserve the efficacy of therapies designed to restore vision.

**Significance Statement:** Retinitis Pigmentosa (RP) is an inherited degenerative disease that affects more than two million people worldwide. RP patients first lose peripheral and low-light vision due to the progressive death of their highly sensitive rod photoreceptors. Photoreceptor degeneration induces pathological noise within the retinal circuit, leading to dramatic structural changes that may hamper therapies to restore visual sensitivity. We discovered a pharmacological treatment that reduces pathological activity in a mouse model of RP without diminishing signaling in surviving circuitry. Partially replacing the neurotransmitter lost when photoreceptors die reduced noise in the retinal circuit without eliminating light sensitivity. This approach could limit the impact of the disease on retinal neurons and preserve the efficacy of subsequent restorative therapies.

## Introduction

Retinitis Pigmentosa (RP) is a prevalent hereditary disease in which photoreceptors degenerate, diminishing retinal sensitivity to visual stimuli (Hamel, 2006; Kuehlewein et al., 2021; O’Neal & Luther, 2023). As degeneration progresses, the inner retinal circuitry exhibits reduced light responses and increased noise levels that compromise visual signal fidelity. The *rd10* mouse model recapitulates a human form of RP in which a missense mutation in the catalytic phosphodiesterase-6 β subunit (PDE6β) disrupts rod photoreceptor function and causes progressive rod death beginning at ages P16-18 (Barhoum et al., 2008; Hamel, 2006; Narayan et al., 2016; O’Neal & Luther, 2023; Stasheff et al., 2011; Yu et al., 2020), followed closely by cone degeneration (Barone et al., 2014; Narayan et al., 2016; Punzo et al., 2009). In both human and mouse, RP causes anatomical and physiological changes at synapses between rods and rod bipolar cells (RBCs) (Denlinger et al., 2020; Fariss et al., 2000; Karlen et al., 2020; Pfeiffer, Anderson, et al., 2020; Pfeiffer, Marc, et al., 2020; Phillips et al., 2010) (fig. 1). Deprived of glutamatergic input from rods, RBC dendrites deform and postsynaptic sites disassemble (Hamel, 2006; Lee et al., 2021; Puthussery et al., 2009). Reducing downstream effects of rod degeneration may therefore prevent synaptic dysfunction and consequent circuit remodeling (Jones et al., 2012; Stefanov et al., 2020) and perhaps preserve the efficacy of subsequent therapies to restore visual sensitivity (Batabyal et al., 2020; Busskamp et al., 2010; Cross et al., 2022; Fenner et al., 2023; Mahato et al., 2020; Meza-Rios et al., 2020; Simon et al., 2020; Tun et al., 2023; van Wyk & Kleinlogel, 2023).

**Figure 1:**
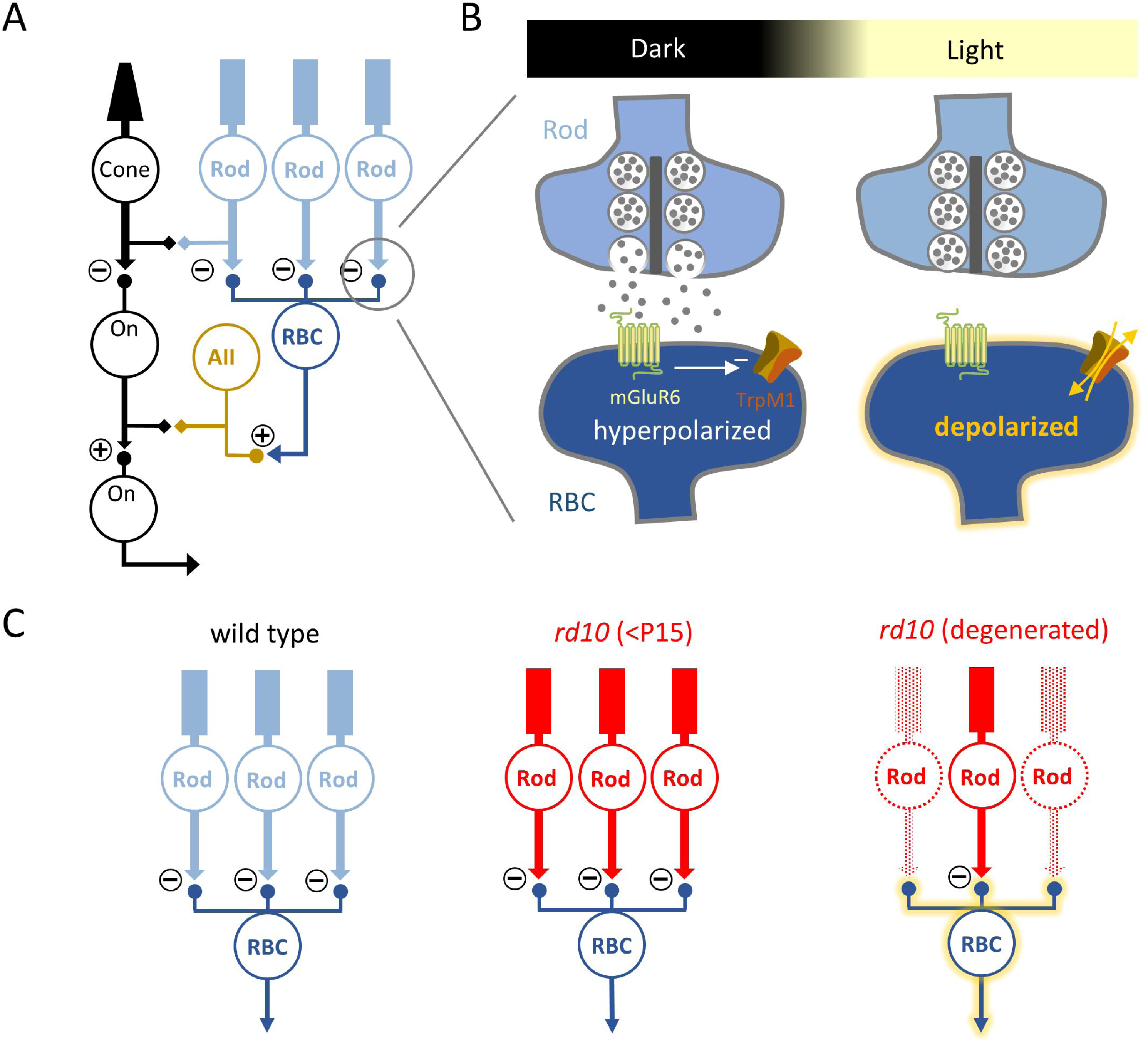
Working model of rod and ON-cone bipolar cell pathways in WT and *rd10* retina. Rod pathway and ON-Cone bipolar cell pathways of WT and *rd10* retina. **(A)** The primary and secondary rod pathway in WT retina. □●, chemical synapses; □□, electrical synapses. **(B)** Schematic showing sign inverting mechanism at the rod-RBC synapse with tonic glutamate release from active rods in the dark keeping RBCs hyperpolarized. In light, rods decrease glutamate release and allow RBCs to depolarize. **(C)** In *rd10* (red), gradual loss of rods (lighter shade) reduces glutamatergic input onto RBCs, resulting in active RBCs lacking rod input in the dark as well as persistent glutamate release from surviving rods due to PDE6β mutation.

In murine models of RP, light-independent circuit noise can be reduced by blocking gap junctions and/or ionotropic glutamate receptors in inner retinal circuitry (Biswas et al., 2014; Busskamp et al., 2010; Cross et al., 2022; Simon et al., 2020; Toychiev et al., 2013; Tun et al., 2023). Since degeneration originates in rods, we hypothesized that targeting type-6 metabotropic glutamate receptors (mGluR6) at rod-RBC synapses might improve signal fidelity in degenerating *rd10* retinas. We first characterized the developmental time course over which rod degeneration in *rd10* retinas degrades visual signaling. We recorded *in vivo* electroretinograms (ERGs) to measure photoreceptor and bipolar cell activity, then measured light-evoked excitatory postsynaptic currents (EPSCs) from ON alpha ganglion cells (ONα RGCs) in an *in vitro* whole-mount retina preparation to evaluate visual signals and noise within the circuit. Rod degeneration reduced light responses and increased circuit noise, effects that were partially ameliorated by mimicking tonic glutamate input to RBCs with the mGluR6 agonist L-2-amino-4-phosphonobutyric acid (L-AP4). A low concentration of L-AP4 reduced noise in ONα RGCs without altering EPSC amplitude. Perforated-patch recordings indicated that RBCs rest at more depolarized potentials in *rd10* retinas, likely reducing the gain of RBC synaptic output to downstream targets (Oesch & Diamond, 2011). Our results indicate that L-AP4 reduces circuit noise and preserves synaptic gain by hyperpolarizing *rd10* RBCs. They suggest a broadly applicable therapy that could ameliorate deleterious downstream effects of photoreceptor degeneration, maintaining visual signaling and potentially preserving the efficacy of subsequent restorative therapies.

## Materials and Methods

### Animals

In all experiments, animals were treated in accordance with National Institutes of Health guidelines, as approved by the National Institute of Neurological Disorders and Stroke Animal Care and Use Committee (ASP-1344). Wild-type (C57Bl6) and *rd10* mice (B6.CXB1-Pde6b^rd10^/J, Jackson Labs) of either sex were used and were housed on a 12:12 light-dark cycle and provided free access to food and water.

### Electroretinograms

*In vivo* ERG responses were recorded following published protocols from the NEI Vision Core (Li et al., 2020) with an Espion E2 Visual Electrophysiology System (Diagnosys, Lowell, MA, USA). Both WT and *rd10* animals were dark adapted overnight (∼10 hours) and anesthetized by i.p. injection of ketamine (100 mg/kg)/xylazine (6 mg/kg) mixture under dim red light. Pupils were dilated with 1% tropicamide and irritation was reduced with 0.5% phenylephrine. Animals were placed on a heating plate maintained at 37°C. Responses were recorded with a gold loop wire electrode placed at the center of the cornea, a reference electrode in the mouth, and a ground electrode in the tail. ERGs were recorded after a further 3 minutes dark adaptation using 20 ms flashes (10^-5^, 10^-4^, 10^-3^, 10^-2^, 1, and 10 cd·s/m^2^).

### Retina dissection for in vitro experiments

Animals were dark-adapted overnight, deeply anaesthetized with isoflurane (Baxter), and euthanized via cervical dislocation. After bilateral enucleation, the eyes were submerged in bicarbonate-buffered Ames medium (∼32°C, Sigma Millipore, 285-295 mOsm) continuously equilibrated with carbogen (95% O_2_/5% CO_2_). The cornea, lens, and iris were removed using small surgical scissors and forceps under a dissecting microscope (Zeiss) and infrared LED illumination (940 nm, ThorLabs), visualized through the microscope eyepieces with infrared image converters (BE Meyers). Retinas were either used immediately or left in their eyecups and stored for up to 6 hours at room temperature in light-proof chambers in carbogen-equilibrated Ames solution.

### Whole-mount retina recordings

To mount the intact retina, the vitreous was removed from the eyecups, and the retina was carefully separated from the pigment epithelium. The retina was mounted photoreceptor-side down on a poly-L-lysine-coated microscope slide (12mm diameter, Corning BioCoat Cellware) that was secured to the bottom of recording chamber with vacuum grease (Dow Corning). The chamber and tissue were superfused continuously in carbogen-equilibrated Ames solution (7-9 mL/min, ∼32°C).

ON αRGCs were targeted based on large soma size and light response characteristics and confirmed morphologically via confocal imaging after recordings. Cell attached and whole cell recordings were made with an electronic amplifier (MultiClamp 700B, Molecular Devices) and signals were digitized (10kHz sample rate; ITC-18, InstruTECH). Voltage-clamp whole-cell recordings were conducted with borosilicate glass electrodes (1.5mm OD, ∼4MΩ) filled with internal solution containing (in mM): 120 CsMeSO_4_, 5 HEPES, 2 NaCl, 2 EGTA, 1MgCl_2_, 6 TEA-Cl, 2 QX-314-Br, 4 Mg-ATP, 0.4 Na-GTP, 14 Tris-phosphocreatine, and 0.1 Alexa 488/Alexa 568 hydrazide (265-270 mOsm, adjusted to pH of 7.4 with CsOH). Absolute voltage values were corrected for a −10 mV liquid junction potential in both cesium- and potassium-based intracellular solutions. L-AP4 (50 nM and 10 µM; Tocris Bioscience) was applied via the bath solution. To isolate excitatory synaptic input, ON αRGCs were held at the estimated reversal potential for excitatory input of −60 mV.

### Retinal slice recordings (paired and perforated patch recordings)

For both whole-cell recordings from RBCs and AII amacrine cells and RBC perforated patch recordings, dissections were performed in room-temperature bicarbonate-based AMES media (285-295 mOsm) equilibrated with carbogen. A section of isolated retinal tissue was embedded in low gelling temperature agarose (3% in HEPES-based AMES media) and then submerged in ice-cold HEPES-based AMES media (285-295 mOsm, NaOH-adjusted to pH 7.4). Transverse slices (200 μm thick) were cut on a vibratome (Leica VT1000S), stored in a light-proof carbogen-equilibrated container, and used for 4-6 hours after slicing. Slices were placed in a recording chamber under a harp (ALA Scientific) and continuously superfused with carbogen-equilibrated Ames medium as above.

Paired recordings from synaptically connected RBCs and AIIs were performed at room temperature (22-25°C) from adult (P28-P90) light adapted C57BL/6 WT and *rd10* mice. Pairs were visually identified using IR-DIC microscopy (Zeiss LSM-510, 40x/1.0 NA objective). Recordings were made in AMES media supplemented with strychnine hydrochloride (3 µM, Sigma Aldrich) and picrotoxin (100 µM, Abcam) to block inhibitory synaptic transmission, and L-AP4 (10 µM, Tocris Bioscience) to minimize synaptic input from other RBCs presynaptic to the recorded AIIs (Singer & Diamond, 2003). Whole-cell voltage-clamp recordings were made from RBCs with pipettes (1.5mm OD borosilicate glass, ∼12 MΩ) filled with solution containing (in mM): 100 CsMeSO4, 10 HEPES, 1.5 BAPTA, 20 TEA-Cl, 4 Mg-ATP, 0.4 Na-GTP, 14 Tris-phosphocreatine (275-280 mOsm, pH = 7.3). For AII voltage-clamp recordings, pipettes (3-6 MΩ) were filled with solution containing (in mM): 95 CsMeSO4, 10 HEPES, 10 EGTA, 20 TEA-Cl, 3 QX314-Br, 4 Mg-ATP, 0.4 Na-GTP, 14 Tris-phosphocreatine (275-280 mOsm, pH 7.3). Both internal solutions were routinely supplemented with fluorescent dyes to confirm the morphology of recorded cells (RBC: 50 µM AlexaFluor 488; AII: 50 µM AlexaFluor 647, both from stock solutions diluted in filtered, deionized water). RBCs were held at different levels of V_pre_ from −79.2 to −29.2 mV in 5 mV increments (conditioning steps) to achieve different equilibrium RRP sizes. From each of these V_pre_ levels we then made a voltage step to −29.2 mV (test step) to complete the release of the RRP (Singer & Diamond, 2006).

For perforated patch recordings, RBCs from P20-25 WT and *rd10* animals, dark-adapted overnight, were identified by their distinct somatic shape and light response characteristics, and confirmed via confocal imaging after breaking into whole cell at the conclusion of perforated patch recordings. Current clamp recordings were conducted using the same glass electrodes as above, filled with internal solution containing (in mM): 125 K-aspartate, 10 KCl, 10 HEPES, 5 N-methyl glucamine-HEDTA, 0.5 CaCl2, 1 ATP-Mg, 0.2 GTP-Mg, and 0.1 Alexa 488/Alexa 594 hydrazide at 265-270 mOsm, adjusted to pH of 7.3 with N-methyl-D-glucamine (NMG)-OH. Beta-Escin (25 μM) was used as a perforating agent. Membrane currents were filtered at 300 Hz and sampled at 10 kHz.

### Visual stimulation and analysis

Full field light stimuli (500 μm diameter spot) were presented using a customized 912 × 1140-pixel digital projector (DLPLCR4500; Texas Instruments) (Franke et al., 2019) driven by a 405 nm LED (ThorLabs) at a frame rate of 60 Hz. Spatial stimuli patterns were created with MATLAB-based software (https://github.com/Schwartz-AlaLaurila-Labs/sa-labs-extension). Photon flux was attenuated to desired levels using a motorized neutral density filter wheel (FW102C, Thorlabs) and routed through the microscope (Scientifica Hyperscope) condenser, which was adjusted so that images were in focus at the plane of the photoreceptor outer segments. Photoisomerization rates were calculated based on a collecting area of 0.85 μm^2^ for rods (Govardovskii et al., 2000; Lyubarsky et al., 2004). Stimuli were centered relative to the recorded cell and focused on the photoreceptor layer. Irradiance (W/m^2^) was converted to photoisomerization rate (R*/rod/s) using the estimated collecting area of rods and cones (0.5 and 0.37 μm^2^, respectively), the 405-nm LED (ThorLabs) emission spectrum, and the photoreceptor absorption spectra (Govardovskii et al., 2000). Light responses were delivered from darkness (to 40R*/rod/s, 1 s step) or from a 500 R*/rod/s background to 505-2500 R*/rod/s (+1 to +400% Weber contrast, 500 ms step).

For voltage clamp experiments, EPSC charge was calculated by integrating the averaged current responses (average of 5 trials) over the stimulus time (1 second) window. Noise (variance) was calculated over the 1 second interval prior to light stimulus; all data for variance and charge calculations were baseline corrected.

Electrophysiological data were analyzed in MATLAB using a custom written open-source package (http://www.github.com/SchwartzNU/SymphonyAnalysis). Figures were constructed in IgorPro 8.04 (Wavemetrics) and Powerpoint.

### Estimating the RRP size

During paired RBC-AII recordings, a step depolarization in the RBC elicited an EPSC in the AII comprising transient and sustained components (Singer & Diamond, 2003). When evoked by large presynaptic voltage steps, the transient component reflects the release of the entire readily-releasable pool (RRP) (Oesch & Diamond, 2011; Singer & Diamond, 2003). AII EPSCs were integrated and a linear fit to the sustained component (50-100 ms after step onset) was extrapolated back to the time that the step was initiated (Neher, 2015; Singer & Diamond, 2006), thereby yielding an accurate measure of the RRP size (Singer and Diamond, 2006).

### Experimental design and statistical analysis

Unless indicated otherwise, normally distributed data are reported as mean ± SD with p-values calculated using 2×3 between subjects ANOVA followed by unpaired two-tailed t-tests for comparisons between WT and *rd10*, and mixed ANOVA followed by paired t-test for comparisons between control and L-AP4 treated retinas. Contrast response curves graphed in figures 5E and 6Gi-iii are shown as median ± upper and lower quartiles, as those data were not normally distributed (Jarque-Bera test). These p-values were determined using permutation and the Wilcoxon Rank Sum test. Resting membrane potentials shown in figure 8C are reported as mean ± SE with p-values calculated using 2×3 between subjects ANOVA followed by unpaired two-tailed t-tests for comparisons between WT and *rd10,* and paired t-test for comparisons between control and L-AP4 treated RBCs.

## Results

### In vivo ERG measurements reveal the time course of retinal dysfunction in rd10 mice

We first characterized the timeline of rod signal degeneration in dark-adapted *rd10* mice (P19-45) using *in vivo* electroretinograms (ERGs). *Rd10* mice typically do not exhibit detectable scotopic (i.e., rod-driven) ERGs after ages P30-35 (Gargini et al., 2007; Jae et al., 2013), although some light sensitivity may remain at later stages of RP (Rodgers et al., 2023; Scalabrino et al., 2023). The a-wave component of the ERG waveform reflects photoreceptor activation, whereas ON bipolar cell responses are contained within the b-wave (Li et al., 2020; Perlman, 1995; Pinto et al., 2007). Both components were smaller in *rd10* mice than in WT even prior to significant rod degeneration (fig. 2A,C,E), likely reflecting the effects of the PDE6β mutation on rod signaling (Bowes et al., 1990; Chang et al., 2007; Gargini et al., 2007; Kuehlewein et al., 2021). At P19, a-waves were not detectable in response to flashes below 0.01 cd·s/m^2^ in WT, and below 0.1 cd·s/m^2^ in *rd10* (fig. 2C). This likely reflects the sensitivity limits of the *in vivo* ERG to detect a-waves, as the larger b-waves were recorded in response to stimuli as dim as 10^-4^ cd·s/m^2^ (fig. 2E) in both WT and *rd10* retinas. In response to brighter flashes, both a- and b-waves in *rd10* retina progressively diminished such that no a-waves were detected by P45 (fig. 2B,D,F), when rods have degenerated completely (Gargini et al., 2007; Puthussery et al., 2009).

**Figure 2:**
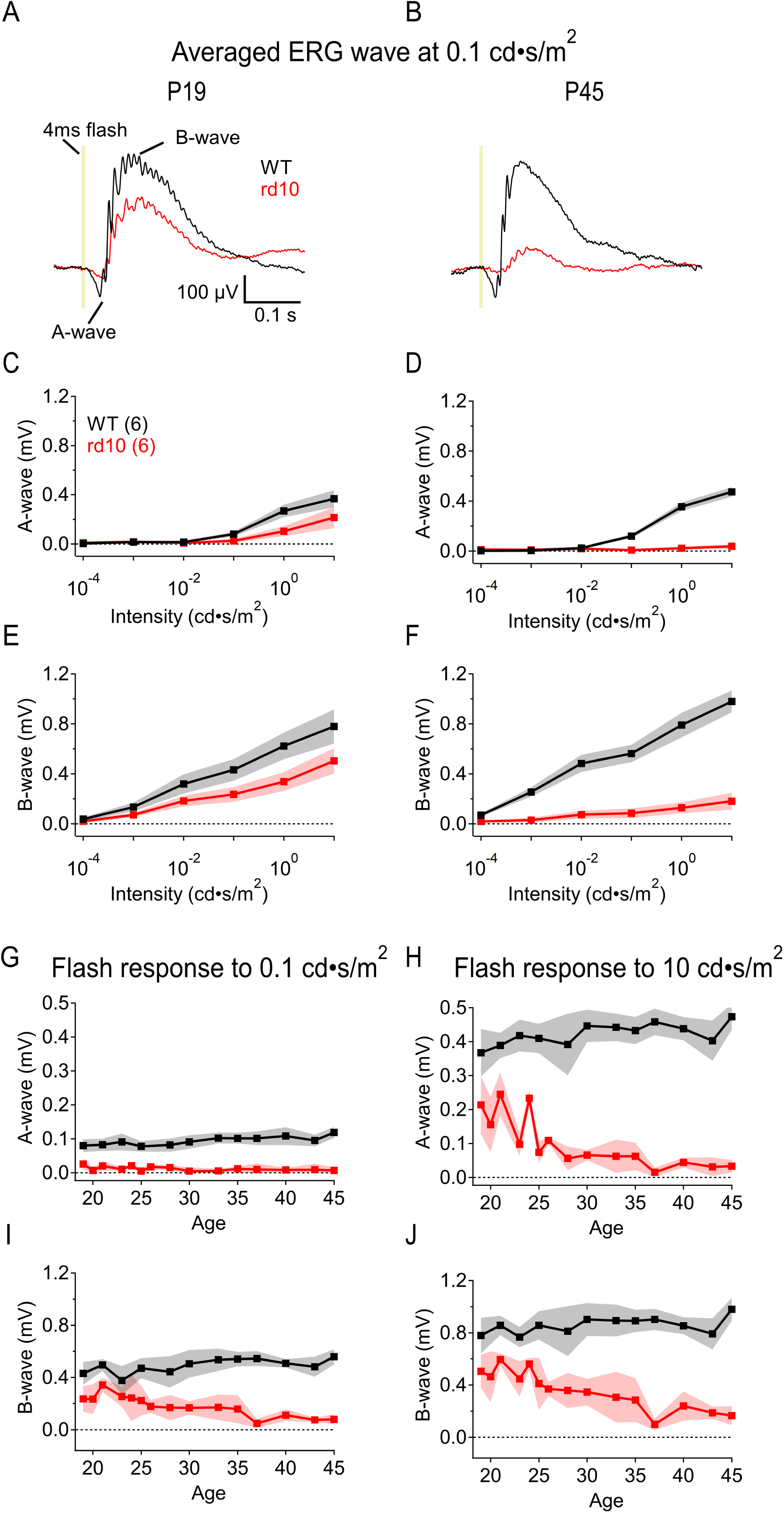
In vivo ERG measurements reveal the time course of retinal dysfunction in rd10 mice. *Rd10* retina *in* vivo ERG responses are diminished after rod degeneration begins. **(A, B)** Example ERGs comparing WT and *rd10* ERGs at P19 (early in *rd10* rod degeneration) and P45 (post *rd10* rod degeneration) respectively. **(C, D)** Summary a-wave amplitudes in WT and *rd10* retina at P19 (C) and P45 (D) (± s.d. (shaded), n = 6 animals/strain) respectively across increasing light stimulus intensities from 10^-4^ to 10 cd·s/m^2^. **(E, F)** Summary b-wave amplitudes in WT and *rd10* retina at P19 (E) and P45 (F) (± s.d. (shaded), n = 6 animals/strain) respectively across increasing light stimulus intensities. **(G, H)** Summary dark-adapted ERG a-wave amplitudes evoked by 0.1 cd·s/m^2^ 20 ms flashes (G) and 10 cd·s/m^2^ flashes (H) from age P19-45 in WT (black) and *rd10* (red) retina. A-waves calculated by peak amplitudes (± s.d. (shaded), n = 8 animals/strain, p < 0.0001 each. **(I, J)** Summary ERG b-wave amplitudes evoked by 0.1 cd·s/m^2^ 20 ms flashes (I) and 10 cd·s/m^2^ flashes (J) from age P19-45 in WT and *rd10* retina. B-waves calculated by peak amplitudes from a-wave peak to b-wave peak (± s.d. (shaded), n = 8 animals/strain; p = 0.025 at 0.1 cd·s/m^2^, p < 0.0001 at 10 cd·s/m^2^.

To examine the age-dependence of signal loss more closely, we delivered a range of flash strengths to P19-45 WT and *rd10* animals (fig. 2G-J). In response to flashes activating only rods (0.1 cd·s/m^2^), or both rods and cones (10 cd·s/m^2^), WT a-wave amplitudes were larger than those in *rd10* across all measured ages (n = 8, p = 0.0001; fig. 2G,H). *Rd10* a-waves could not be resolved at any age in response to dimmer flashes, whereas WT a-wave amplitudes increased with age. In responses to brighter flashes, *rd10* a-wave amplitudes were 38 ± 29% lower than those in WT (P19; n = 8, p < 0.0001; fig. 2H). In contrast to WT, *rd10* a-waves were reduced 60 ± 20% by P25 and were almost undetectable by P37 (fig. 2H). *Rd10* b-wave amplitudes were also significantly lower than in WT and were reduced 65 ± 56% by P35 (0.1 cd·s/m^2^: p = 0.025, n = 8; 10 cd·s/m^2^: p < 0.0001, n = 8; fig. 2I,J). The largest *rd10* b-waves reached only 65% of WT maximum amplitudes (at P19-24), suggesting that the collective RBC activity over the entire age range is less in *rd10* than in WT animals.

### Rod degeneration degrades scotopic signal fidelity in ONα RGCs

To measure physiological circuit output in dark-adapted WT and *rd10* retinas, we recorded from RGCs in the *in vitro* whole-mount retina across the same age range as the *in vivo* experiments. Whereas ERG recordings yield population light responses, whole-cell voltage-clamp recordings of excitatory postsynaptic currents (EPSCs) in RGCs report downstream signals and noise in individual neurons. We focused on sustained ONα RGCs, which have been well characterized physiologically and anatomically (Bleckert et al., 2013; Goetz et al., 2022; Krieger et al., 2017; Laboissonniere et al., 2019). We grouped recorded RGCs into three age ranges: P16-20, the beginning stages of rod degeneration in *rd10*; P21-25, over which time more than 50% of *rd10* rods die; and P26-30, when rod degeneration is mostly complete (Barhoum et al., 2008; Jae et al., 2013; Kim et al., 2018; Li et al., 2018; Puthussery et al., 2009). We first recorded synaptic dark noise and light responses to 1 s light steps (40R*/rod/s; fig. 3A). Cells were filled through the patch pipette with Alexa 488 to identify any morphological differences between WT and *rd10* RGCs, or between age groups; no qualitative differences were observed across ages or between WT and *rd10* (fig. 3B).

**Figure 3:**
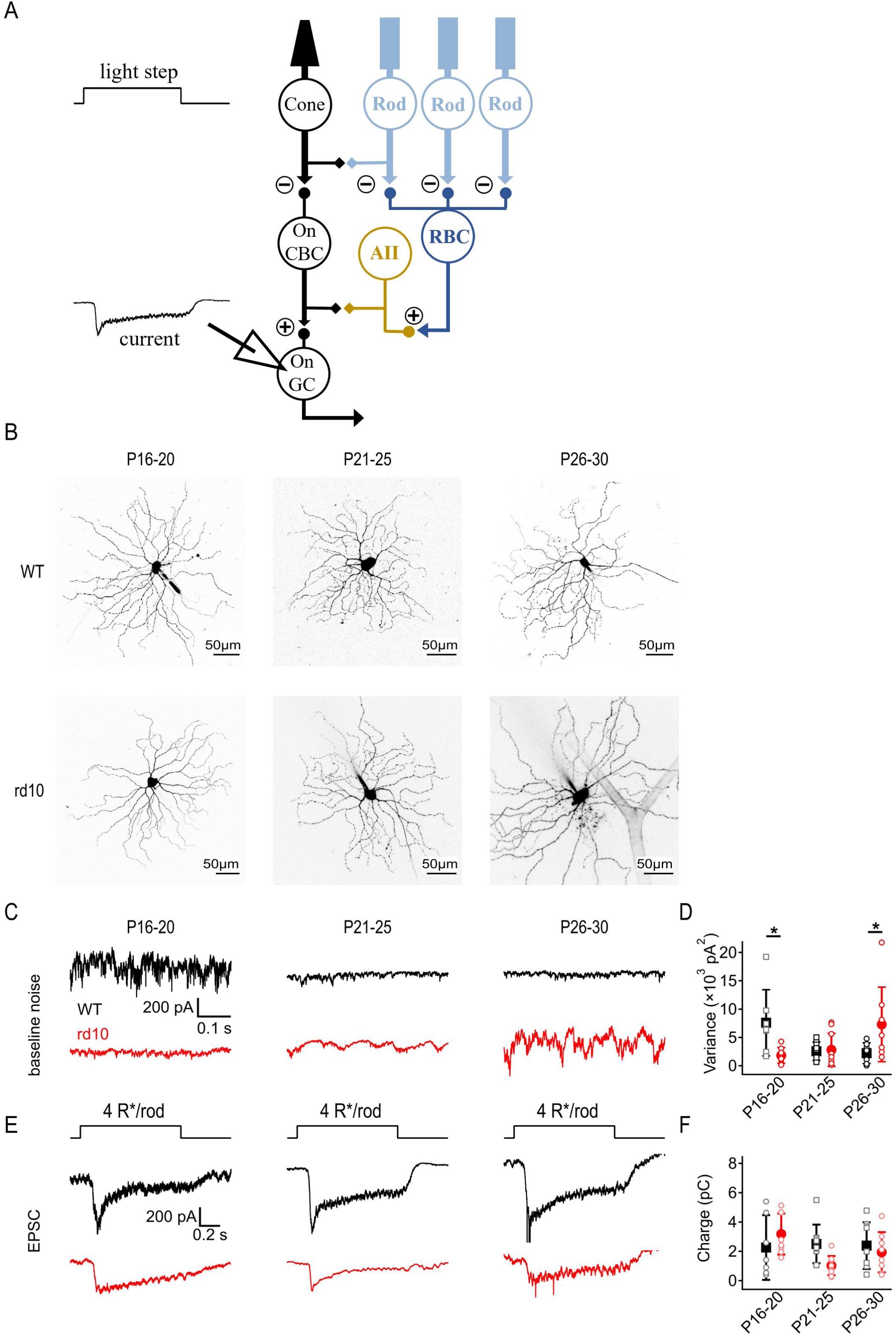
Rod degeneration degrades scotopic signal fidelity in ONα RGCs. Dark-adapted *rd10* ONα RGCs become increasingly noisy after rod degeneration begins, while WT ONα RGCs are noisy during earlier stages of development but become quieter as the retina matures. **(A)** Excitatory post synaptic currents were recorded from voltage clamp whole cell recordings of ONα RGCs. Light stimulus: 1s step of 40R*/rod/s light from darkness. **(B)** Examples of Alexa 488 filled WT and *rd10* RGCs at each age group imaged following recordings. **(C)** Example traces of baseline EPSCs from WT and *rd10* ONα RGCs at 3 age groups showing noise during early rod degeneration (P16-20), peak rod degeneration (P21-25), and end of rod degeneration (P26-30). **(D)** Summary of baseline variance of WT and *rd10* ONα RGCs at 3 age groups (± s.d., n = 10 cells/age/strain) showing noise levels during early rod degeneration (P16-20), peak rod degeneration (P21-25), and end of rod degeneration (P26-30). WTs are significantly noisier than *rd10*s at the youngest (P16-20, p = 0.015,) and oldest ages (P26-30, p = 0.017). **(E)** Example EPSCs evoked by 40R*/rod/s light steps WT and *rd10* ONα RGCs during early rod degeneration (P16-20), peak rod degeneration (P21-25), and end of rod degeneration (P26-30). **(F)** Summary of EPSC charge transfer in WT and *rd10* ONα RGCs at all 3 age groups. No significant changes were observed between WT or rd10 ONα RGC response amplitudes across age groups.

WT ONα RGCs exhibited synaptic dark noise that decreased with age (fig. 3C,D). By contrast, dark noise was relatively low in P16-20 *rd10* RGCs and increased as rod degeneration progressed (fig. 3C,D), consistent with previous work showing spontaneous noise and oscillations in *rd10* retinas (Biswas et al., 2014; Goo et al., 2015; Trenholm & Awatramani, 2015). We observed increasing variability in noise levels between *rd10* RGCs with age (fig. 3D). Rod-driven EPSCs recorded from *rd10* ONα RGCs steadily decreased as degeneration progressed, while EPSCs in WT ONα RGCs were similar across all ages (fig. 3E,F).

### Low dose of L-AP4 suppresses circuit noise under scotopic conditions

The results thus far indicate that *rd10* rod degeneration decreases visual responses and increases circuit noise, degrading visual signal fidelity. We hypothesized that a low dose of mGluR6 agonist L-AP4 (binding affinity ∼2 μM) (Naples & Hampson, 2001; Thomsen, 1997) might stabilize RBC activity without completely blocking responses to rod input, thereby enhancing visual signaling in *rd10* animals. Consistent with this prediction, 50 nM L-AP4 lowered dark noise in both WT and *rd10* ONα RGCs across all age groups (fig. 4A-C). The effect of 50 nM L-AP4 on ONα RGC EPSC amplitudes was highly variable between cells in both WT and *rd10*, leading to an insignificant effect overall (fig. 4D-F). Taken together, these results suggest that a low dose of L-AP4 improves signal-to-noise characteristics in the degenerating retina.

**Figure 4:**
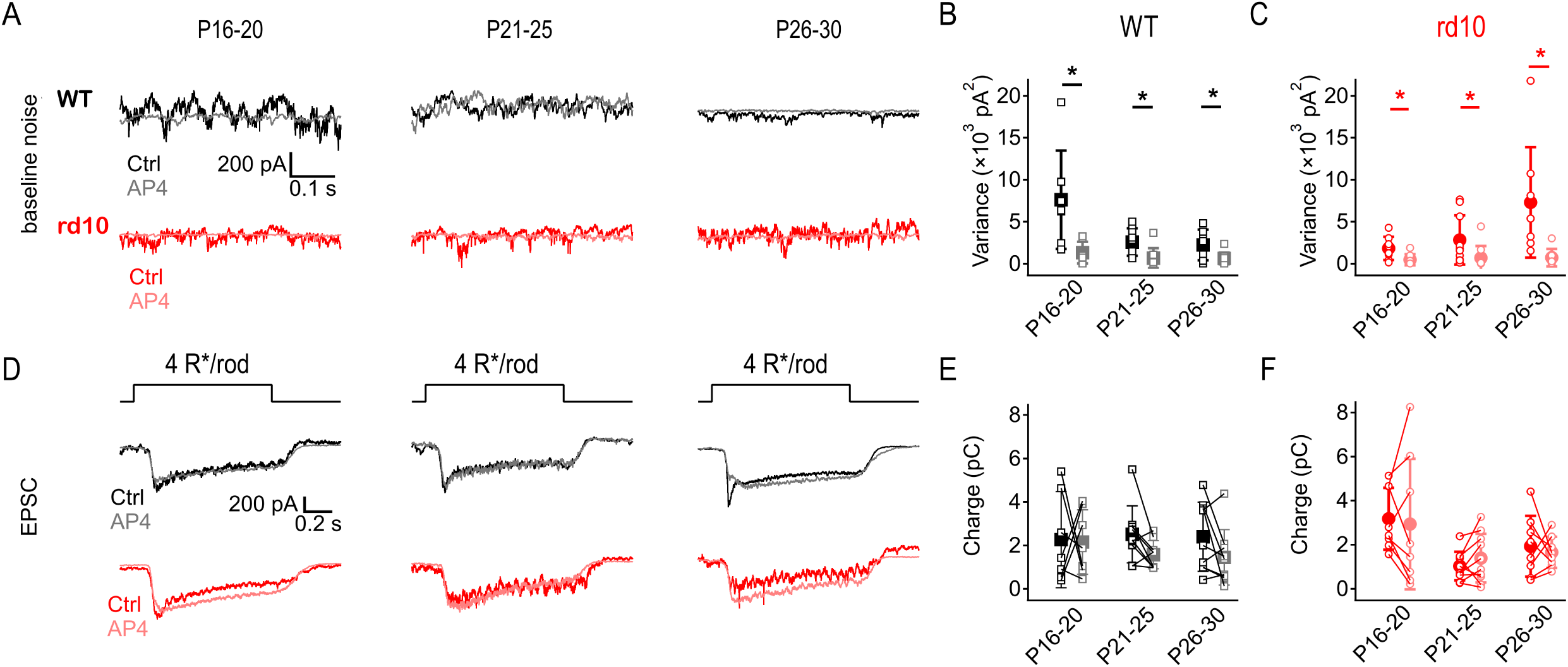
Low dose of L-AP4 suppresses circuit noise under scotopic conditions. Low-dose of L-AP4 reduced noise but did not affect light responses under dark-adapted conditions. **(A)** Example traces of baseline recordings of WT and *rd10* ONα RGCs at 3 age ranges showing noise during early rod degeneration (P16-20), peak rod degeneration (P21-25), and end of rod degeneration (P26-30) before and after application of 50 nM L-AP4 (lighter shades). L-AP4 reduced noise in all age groups in both strains. **(B, C)** Summary of baseline noise in WT (B) and *rd10* (C) ONα RGCs at 3 age groups (± s.d., n = 10 cells/age/strain) showing noise at each age range before (x-axis) and after application of 50 nM L-AP4 (y-axis). L-AP4 reduced noise in all ages of both WT (p = 0.024, 0.015, and 0.022) and *rd10* RGCs (p = 0.048, 0.02, and 0.042 respectively in increasing age groups). **(D)** Example EPSCs evoked by 40R*/rod/s light steps in WT and *rd10* ONα RGCs during early rod degeneration (P16-20), peak rod degeneration (P21-25), and end of rod degeneration (P26-30) before and after application of 50 nM L-AP4 (lighter shades). **(E, F)** Summary of EPSC charge transfer in WT (E) and *rd10* (F) ONα RGCs at all 3 age groups before and after application of 50 nM L-AP4 (lighter shades). No significant changes observed with L-AP4 in either WT or rd10 ONα RGC response amplitudes within each age group.

### Rod degeneration degrades mesopic contrast signals in ONα RGCs

In the experiments presented thus far, dim light stimuli were delivered from darkness to evoke primarily rod-mediated responses. Even at night, however, animals navigating their visual world must distinguish the relative luminance of objects from their immediate surroundings, conditions that are best described in terms of contrast. To examine the effects of rod degeneration on contrast encoding, we delivered stimuli upon a 500 R*/rod/s full-field background, mesopic conditions under which contrast responses in RGCs are mediated by both the rod and cone pathways (Nath et al., 2023). Similar to our results under scotopic conditions (fig. 3C,D), WT ONα RGCs exhibited background noise at P16-20 that decreased with age (fig. 5A,B). *Rd10* ONα RGCs exhibited the opposite trend, with low noise levels early in development that increased significantly by P26-30 (p = 0.046; fig. 5B), likely in response to severe rod loss (Barhoum et al., 2008; Pennesi et al., 2012; Puthussery et al., 2009).

**Figure 5:**
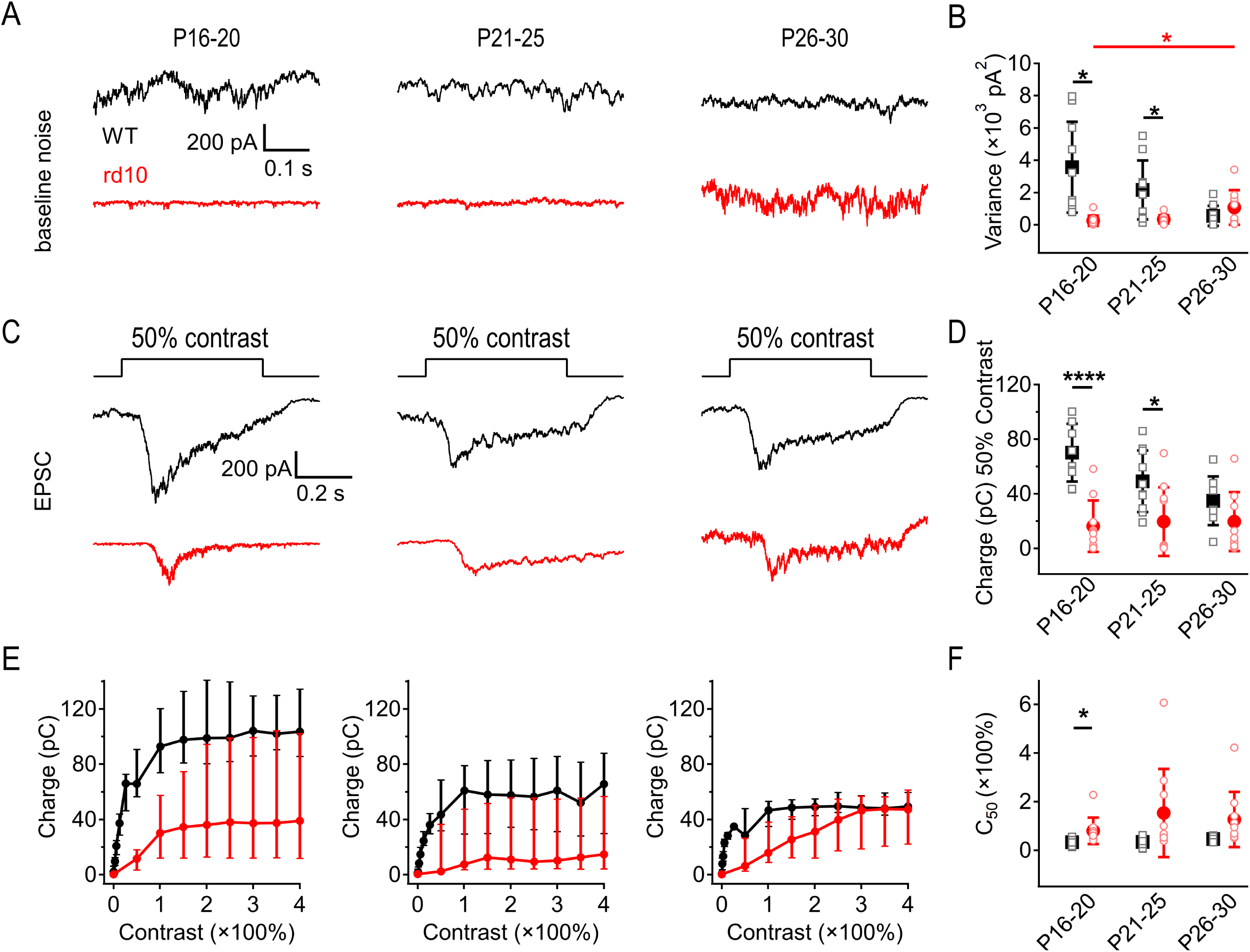
Rod degeneration degrades mesopic contrast signals in ONα RGCs. *Rd10* ONα RGCs under mesopic luminance become noisy only after rod degeneration begins, while WT ONα RGCs are noisy during development and become quieter as the retina matures. **(A)** Example traces of voltage clamp whole cell recordings of WT and *rd10* ONα RGCs at three age groups showing baseline noise during early rod degeneration (P16-20), peak rod degeneration (P21-25), and the end of rod degeneration (P26-30) under mesopic (500 R*/rod/s) luminance. **(B)** Summary of WT and *rd10* ONα RGC noise across three age groups (± s.d., n = 10 cells/age/strain). *Rd10* RGCs exhibited relatively low spontaneous activity at P16-20 but became significantly noisier following rod degeneration (P26-30, p = 0.046). WT RGCs exhibited higher noise at both P16-20 (p = 0.014) and P21-25 (p = 0.034). **(C)** Example responses to +50% contrast step measured under voltage clamp (V_hold_ = −70 mV) in WT and *rd10* ONα RGCs during early rod degeneration (P16-20), peak rod degeneration (P21-25), and end of rod degeneration (P26-30). **(D)** Summary of EPSC charge transfer evoked by +50% contrast steps in WT and *rd10* ONα RGCs at all 3 age groups under mesopic (500 R*/rod/s) background luminance. WT ONα RGC responses were larger than those of *rd10*s in the two younger age groups (± s.d., n = 10 cells/age/strain, p < 0.0001 and p = 0.039). **(E)** ONα RGC responses of WT and *rd10* retina across contrasts up to +400% on 500R*/rod/s background. WT ONα RGCs display larger responses than *rd10*s in the youngest age group (P16-20: p = 0.003). WT ONα RGCs reached saturation at +100% to +150% contrast, but *rd10* RGC responses did not saturate until +250% to +300% contrast (median ± upper and lower quartile, n = 10 cells/age/strain). **(F)** WT ONα RGCs consistently reach half maximum light response at 50% contrast across ages. However, *rd10* ONα RGCs reach half maximal contrast increased with age from +80% to +150% (± s.d., n = 10 cells/age/strain; P16-20 C_50_ p = 0.024).

In *rd10* ONα RGCs, EPSCs evoked by 750 R*/rod/s stimuli (+50% contrast) were 45% - 85% smaller than WT EPSCs across all ages and exhibited increased noise throughout the light response (fig. 5C,D). ONα RGC responses across a range of contrast stimuli (up to +400%) were significantly larger in WT compared to *rd10* in the youngest age group (P16-20: p = 0.003; fig. 5E). *Rd10* ONα RGCs exhibited much greater variability across cells in EPSC amplitude compared to WT (fig. 5E), an effect that may arise from variability in the extent of degeneration across individual *rd10* retinas (Barhoum et al., 2008; Puthussery et al., 2009; Puthussery & Taylor, 2010). WT ONα RGCs produced a half-maximal light response (C_50_) at an average contrast of 50 ± 7% (fig. 5F), consistent with previous reports (Grimes et al., 2014; Schwartz et al., 2012). *Rd10* ONα RGCs exhibited higher C_50_ values (C_50_ = 120 ± 37%) and were less responsive than WT to lower contrast stimuli across all ages (fig. 5F). These results indicate that rod degeneration in *rd10* retinas degrades the fidelity of mesopic contrast signals.

### A low dose of L-AP4 decreases mesopic synaptic noise in ONα RGCs

The mGluR6 agonist, L-AP4, acts on ON cone bipolar cells as well as RBCs and may therefore affect noise and light responses in the cone pathway under mesopic conditions. We found that 50 nM L-AP4 had no significant effect on noise in WT ONα RGCs (fig. 6A,B), but reduced noise in the two older age ranges in *rd10* ONα RGCs (P21-25: 56 ± 25%, p = 0.024, P26-30: 59 ± 23%, p = 0.05) (fig. 6A,C). EPSCs evoked by 750 R*/rod/s stimuli (+50% contrast steps) were not significantly altered by 50 nM L-AP4 in either WT (fig. 6D,E) or *rd10* ONα RGCs (fig. 6D,F).

**Figure 6:**
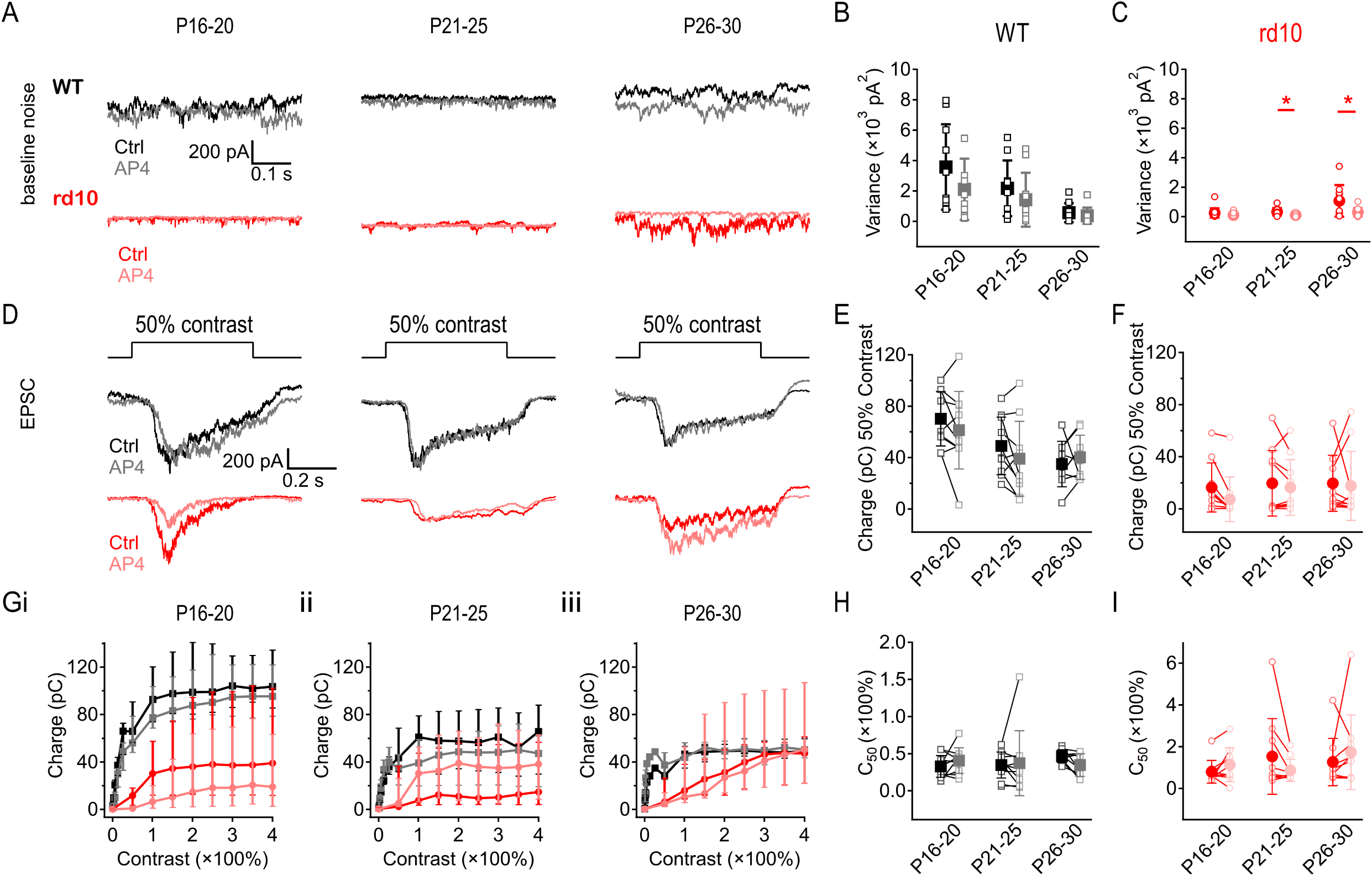
A low dose of L-AP4 decreases mesopic synaptic noise in ONα RGCs. 50 nM L-AP4 reduced noise under mesopic background luminance but did not affect contrast responses. **(A)** Example traces of baseline EPSCs (V_hold_ = −70 mV) in WT and *rd10* ONα RGCs during early rod degeneration (P16-20), peak rod degeneration (P21-25), and end of rod degeneration (P26-30) before and after application of 50 nM L-AP4 (lighter shades). L-AP4 reduced noise in all age groups in both WT and *rd10* RGCs. **(B, C)** Summary of baseline noise in WT (B) and *rd10* (C) ONα RGCs at 3 age groups (± s.d., n = 10 cells/age/strain) showing noise at each age range before (darker shades) and after application of 50nM L-AP4 (lighter shades). L-AP4 did not change noise in WT at any age, but reduced noise in older ages of *rd10* (P21-25, p = 0.024; P26-30, p = 0.05). **(D)** Example EPSCs evoked by +50% contrast steps in WT and *rd10* ONα RGCs during early rod degeneration (P16-20), peak rod degeneration (P21-25), and end of rod degeneration (P26-30) before (darker shades) and after application of 50 nM L-AP4 (lighter shades). **(E, F)** Summary of EPSC charge transfer evoked by +50% contrast steps in WT (E) and *rd10* (F) ONα RGCs at all 3 age groups before and after application of 50nM L-AP4 (lighter shades). No significant changes were observed with L-AP4. **(G)** ONα RGC responses of WT and *rd10* retinas (median ± upper and lower quartile, n = 10 cells/age/strain) across contrasts from 0 to +400% upon a 500R* background before (darker shades) and after application of 50 nM L-AP4 (lighter shades). L-AP4 reduced the response amplitude of P16-20 *rd10* RGCs (p ≤ 0) but did not significantly alter light response amplitudes at P21-25 or P26-30. L-AP4 did not affect response amplitudes of WT RGC contrast responses at any age. **(H, I)** Half maximum light responses of WT (H) and *rd10* (I) ONα RGCs to contrast steps before and after application of 50 nM L-AP4 (lighter shades). WT reached half maximum light response at around 50% contrast, whereas *rd10* reached half max at around 100% contrast (± s.d., n = 10 cells/age/strain).

Lastly, we assessed the effects of 50 nM L-AP4 on responses to positive contrast steps up to +400% contrast (2500R*/rod/s) with L-AP4 (fig. 6G). L-AP4 reduced EPSC amplitudes in *rd10* P16-20 RGCs from 50% - 400% contrast stimuli (p = 0.0002, fig. 6Gi) but had no significant effect on contrast responses in the older age groups, nor any effect on WT contrast response amplitudes at any age (fig. 6Gi-iii). L-AP4 slightly reduced the half maximum contrast sensitivity of P26-30 WT ONα RGCs from 47 ± 8% contrast to 40 ± 15% contrast (p = 0.014) but did not change contrast sensitivity at other ages (fig. 6H). *Rd10* RGCs showed no significant change in contrast sensitivity with low dose L-AP4 regardless of age (fig. 6I). L-AP4 may not have affected mGluR6 synapses equally, and differing degrees of degeneration across *rd10* rods likely contributed to the variability of L-AP4’s effect on signal, but the significant decrease seen in *rd10* RGC noise with L-AP4 was consistent (fig. 6A,G). These results suggest that low-dose L-AP4 in *rd10* retinas reduced noise without altering signal under mesopic luminance, thus improving retinal encoding even as rods continued to degenerate. This change was most evident in *rd10* animals age P21-30 before the complete loss of rods.

### Rd10 retina does not tolerate intravitreal injections without photoreceptor damage

To test whether L-AP4 could improve visual function over the time course of degeneration in RP, we intraocularly injected L-AP4 into *rd10* mouse eyes with the intent of recording *in vivo* ERGs over the course of rod degeneration. L-AP4 injection (10μM, 0.5 µL) did not affect a-waves in WT animals (n = 6; pre-injection vs. one day post-injection) and reduced b-waves evoked by 10 cd·s/m^2^ flashes by 40 ± 25% (p < 0.0001; fig. 7Ai-ii), an expected result from a saturating concentration of L-AP4. In *rd10* animals, however, L-AP4 injection reduced a-waves by 70 ± 28% (p = 0.0008; fig. 7Bi). Similar results were observed upon injection of normal saline (p = 0.002; fig. 7Bi), indicating that intraocular injections were poorly tolerated by *rd10* photoreceptors, preventing further study of the longitudinal effects of L-AP4 on visual signaling in *rd10* retinas using intraocular injections.

**Figure 7:**
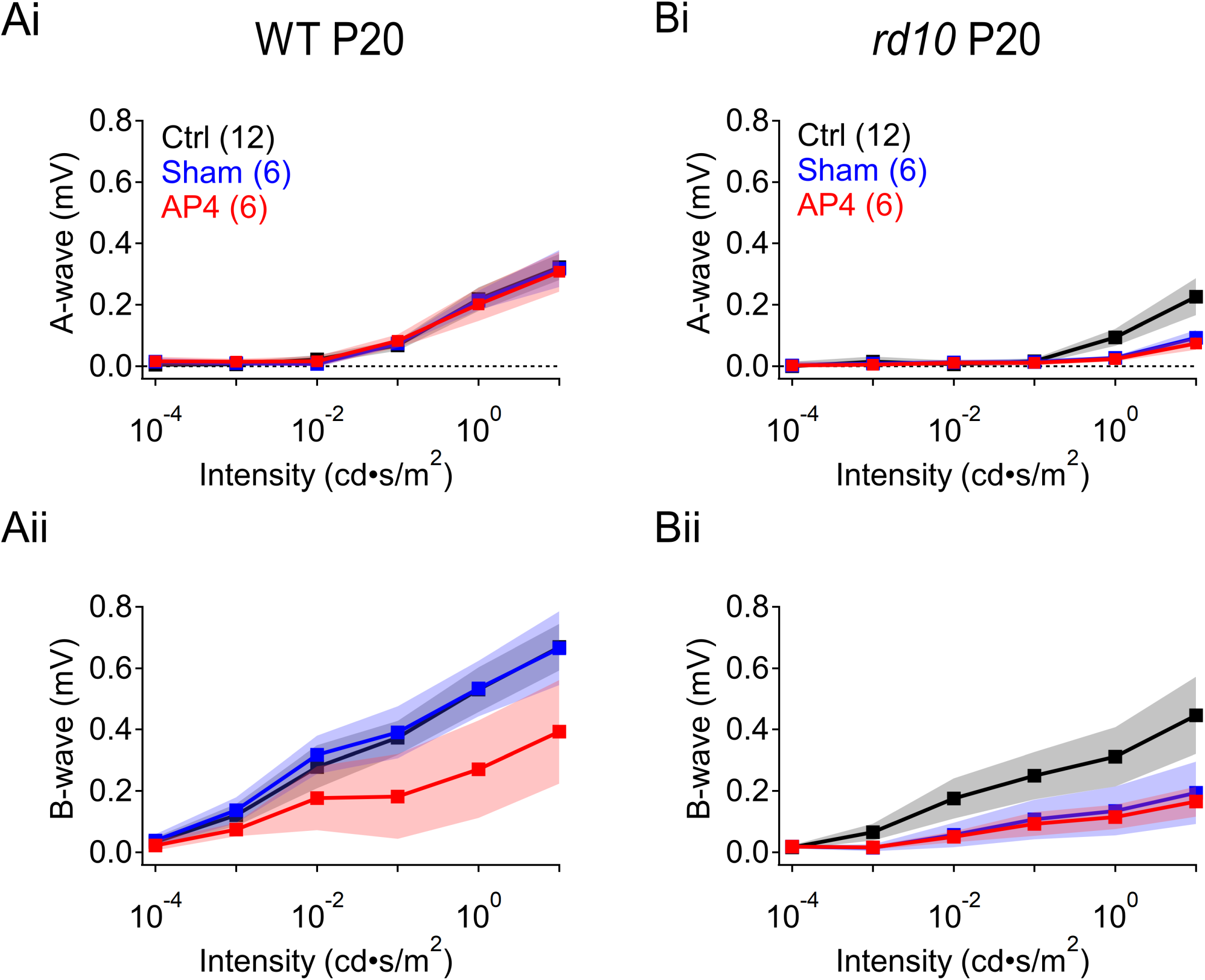
Rd10 retina does not tolerate intravitreal injections without photoreceptor damage. *Rd10* retina did not withstand intraocular injections. **(A)** *In vivo* ERG recordings of WT animals at age P20 pre-injection (black); post NS injection (blue); and post 1mM L-AP4 injection (red). (**Ai**) Quantified a-wave amplitudes at increasing light flash intensities from 10^-4^ to 10 cd·s/m^2^. (**Aii**) Quantified b-wave amplitudes at increasing light flash intensities from 10^-4^ to 10 cd·s/m^2^ (± s.d., n = 6 animals/strain). **(B)** *In vivo* ERG recordings of *rd10* animals at age P20 pre-injection (black); post NS injection (blue); and post 10 mM L-AP4 injection (red). (**Bi**) Quantified a-wave amplitudes at increasing light flash intensities from 10^-4^ to 10 cd·s/m^2^. (**Bii**) Quantified b-wave amplitudes at increasing light flash intensities from 10^-4^ to 10 cd·s/m^2^ (± s.d., n = 6 animals/strain).

### RBCs are more depolarized at rest in rd10 retinas

Experiments presented thus far show that a low dose of L-AP4 during *rd10* rod degeneration reduced circuit noise but not light-evoked responses in RGCs. To dissect the mechanisms by which L-AP4 influenced circuit noise properties, we first measured the resting membrane potential (RMP) directly from P20-25 RBCs with perforated patch recordings (fig. 8A). We hypothesized that at this intermediate stage of degeneration, progressive rod loss and consequent loss of glutamatergic input would depolarize the RMP of RBCs. Indeed, *rd10* RBCs rested significantly more depolarized (RMP = −43 ± 2 mV, n = 8) compared to WT RBCs (RMP = −53 ± 1 mV, n = 8, p = 0.003); (Dunn & Rieke, 2006; Grimes et al., 2014; Herrmann et al., 2011; Oltedal et al., 2009). A saturating concentration of L-AP4 (10 μM) re-hyperpolarized the RMP of *rd10* RBC to −52 ± 2 mV (p = 0.026, n = 8), closer to that of WT RBCs (fig. 8B,C). 50 nM L-AP4, which reduced noise levels in RGCs (fig. 4,6), exerted variable effects on the RMP of individual RBCs that were overall insignificant. These results suggest that recordings from downstream RGCs, which collect input mediated by many RBCs, provide a more sensitive indication of the subtle effects of 50 nM L-AP4 on circuit noise. As a result of photoreceptor death, postsynaptic changes in the *rd10* glutamatergic signaling cascade occur where mGluR6 (Barhoum et al., 2008; Gargini et al., 2007; Puthussery et al., 2009) and TRPM1 (Gayet-Primo & Puthussery, 2015) both lose dendritic synaptic expression but retain somatic expression. This glutamatergic signaling reorganization, in addition to a loss of glutamate from degenerated rods, may also contribute to the depolarized RMP observed in our data.

**Figure 8:**
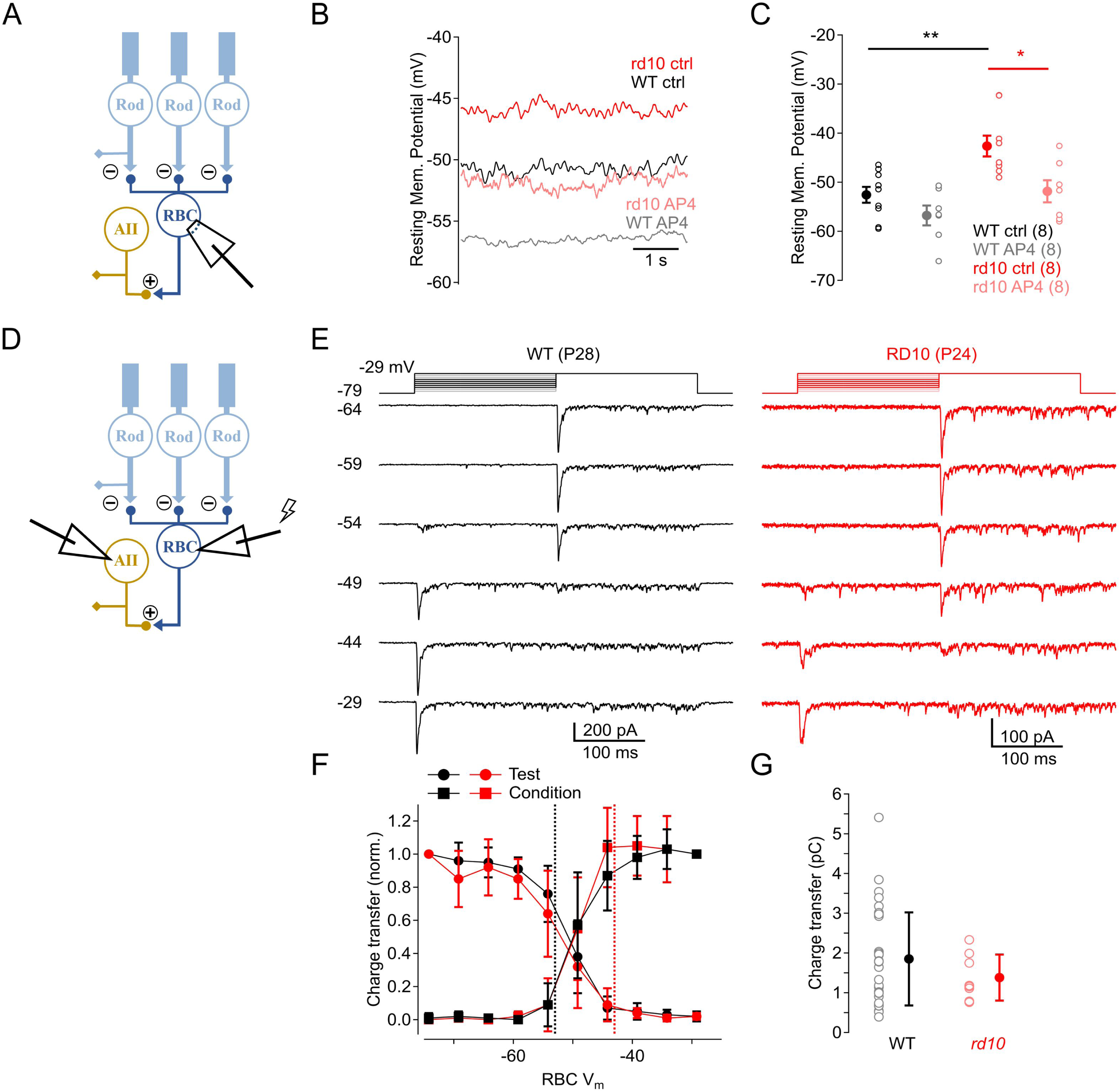
RBCs are more depolarized at rest in rd10 retinas. *Rd10* RBCs rest more depolarized than WT RBCs and reduce synaptic output during rod degeneration. L-AP4 hyperpolarizes the depolarized *rd10* RBCs closer to the resting potential of WT RBCs. **(A)** Schematic showing experimental setup of perforated patch recordings of RBCs. **(B)** Raw trace of resting membrane potential (RMP) in a *rd10* RBC in control (red) and in the presence of 10 μM L-AP4 (pink), and a WT RBC in control (black) and in the presence of 10 μM L-AP4 (gray). **(C)** Summary of RMP in *rd10* RBCs in control (red), and in the presence of 10 μM L-AP4 (pink), and RMP in WT RBC in control (black), and in the presence of 10 μM L-AP4 (gray) (± s.e., n = 8 cells/strain). **(D)** Schematic showing experimental setup of voltage clamp paired RBC-AII recordings. **(E)** Example trace showing WT (black) and *rd10* (red) RBC held at different levels of V_cond_ from −79.2 to −29.2 mV in 5 mV increments (conditioning steps) to achieve different equilibrium RRP sizes. From each of these V_cond_ levels followed a voltage step to −29.2 mV (test step) to complete the release of the RRP, and recorded voltage evoked EPSCs from AIIs. **(F)** Average normalized integrated current (± s.d., WT n = 35, *rd10* n = 8 cells) measured over transient response to V_cond_ (squares) and V_test_ (circles) steps for both WT (black) and *rd10* (red) retina pairs. Currents were normalized to the largest test step without a conditioning step. Vertical dash lines indicate average RMP of WT (black) and *rd10* (red) RBCs. **(G)** Summary of EPSC charge. Data from individual cell pairs in WT (black) and *rd10* (red) are shown in open circles; averaged data shown in closed circles (± s.d., WT n = 35, *rd10* n = 8 cells).

### Synaptic output from the RBC is reduced in early stages of retinal degeneration

The size of the readily-releasable vesicle pool (RRP) at RBC synapses reflects a balance between vesicle release and replenishment that depends on RBC membrane potential (V_RBC_) (Graydon et al., 2018; Oesch & Diamond, 2011). As V_RBC_ depolarizes in response to increased ambient luminance, ongoing RBC release also increases, thereby reducing the number of vesicles available for release and, consequently, the synaptic gain in response to subsequent visual stimuli. These findings suggest that L-AP4, by hyperpolarizing the RMP in *rd10* RBCs, may increase the synaptic gain between RBCs and postsynaptic AII amacrine cells. This could explain, at least in part, how L-AP4 reduces noise in RGCs without significantly decreasing EPSC amplitude (fig. 6). This hypothesis presumes that the release characteristics at RBC-AII synapses in *rd10* retinas are similar to those in WT. To test this, we made dual whole-cell voltage clamp recordings from synaptically coupled RBC-AII pairs in acute retinal slices (Singer & Diamond, 2003); fig. 8D-G). Voltage steps delivered to the presynaptic RBC elicited EPSCs in the postsynaptic AII (Figure 8E). A step to −29 mV elicited a large EPSC, the transient component of which reflects release of the entire RRP (Singer & Diamond, 2006). Smaller steps evoked release and a consequent decrease in the response to a subsequent large step, demonstrating how V_RBC_ influences the size of the RRP (Oesch & Diamond, 2011) (fig. 8F).

EPSCs elicited by large steps were 24% smaller in *rd10* pairs, although variability between pairs rendered this effect statistically insignificant (fig. 8G). Importantly, the dependence of V_RBC_ on RRP size in *rd10* pairs was similar to that observed in WT (Figure 8F): The size of the RRP varied steeply at about −50 mV, near the RMP in WT RBCs, suggesting that the more depolarized RMP observed in *rd10* RBCs would significantly decrease the RRP and, consequently, the synaptic gain of RBC-AII synapses – an effect that would be partially ameliorated by L-AP4 (Figure 8C).

## Discussion

The results presented here show that L-AP4 reduced RP-induced noise in *rd10* circuitry without compromising light-evoked responses in ONα RGCs, thereby improving visual signal fidelity even as photoreceptors degenerate. These data suggest L-AP4 as a potential therapy during photoreceptor degeneration to replace glutamatergic input to ON bipolar cells that is lost when photoreceptors die. During rod degeneration, *rd10* RBCs rest at more depolarized levels than in WT, likely reducing the gain of their synaptic output to AIIs (fig. 8). By re-hyperpolarizing *rd10* RBCs, a low dose of L-AP4 reduced circuit noise and restored synaptic gain. This novel approach should work in retinas expressing any RP-causing mutation and could complement or precede other therapies, such as prosthetic photoreceptors <u>(B.-Y. Wang et al., 2022)</u> or gene therapy (Bennett et al., 1996; Pang et al., 2008; Scalabrino et al., 2023; Tun et al., 2023), to restore vision lost to this disease. Stabilizing visual signaling in the inner retina during photoreceptor degeneration also may delay or ameliorate circuit remodeling (T. Wang et al., 2019) and thereby preserve the efficacy of subsequent restorative therapies.

### Effects of degeneration vary within and between individual retinas

Photoreceptor degeneration in RP progresses at varying rates depending on the mutation (S. Daiger et al., 2013; S. P. Daiger et al., 2007): In humans, some RP patients present visual deficits by 10 years of age (Y. N. Kim et al., 2020), whereas others do not exhibit symptoms until middle age (Kuehlewein et al., 2021; McLaughlin et al., 1995). In the *rd10* mouse model, rod degeneration begins at ∼P16 in the central retina and progresses gradually to the periphery (Gargini et al., 2007). Single-cell and population recordings indicate that the physiological impact of disease progression varies between neighboring regions of the same retina (Puthussery et al., 2009; Stasheff et al., 2011). We also observed high cell-cell variability in noise levels and light responses in *rd10* RGC recordings (fig. 3-6), likely due to the variable impact of rod degeneration on individual upstream RBCs.

### Reducing noise in the primary rod pathway could enhance cone-mediated vision

The rod and cone pathways interact at several points within the retinal circuit. Rods contact cones directly through gap junctions (Schneeweis & Schnapf, 1995), and AII amacrine cells relay rod pathway signals to cone bipolar cell terminals (Völgyi et al., 2004). In low light, the high-gain primary rod pathway amplifies both signals and noise; as luminance levels increase, the gain of this pathway is reduced, clearing the way for smaller cone pathway signals to pass through. Rod degeneration in *rd10* increases noise levels in the electrically-coupled ON cone bipolar cell to AII network (Goo et al., 2015; Jae et al., 2013; Stasheff et al., 2011; Trenholm & Awatramani, 2015), effects that are eliminated by blocking ionotropic glutamate receptors and gap junctions (Biswas et al., 2014). Limiting pathological signals in RBCs would reduce noise in AIIs, thereby preventing contamination of cone pathway signals. By decreasing noise without reducing light responses, the low-dose L-AP4 treatment described here may improve visual signaling in degenerating retinas across light levels.

### A low dose of L-AP4 reduces noise while preserving signal

L-AP4 (EC_50_ ∼ 2 μM; (Naples & Hampson, 2001; Thomsen, 1997) is typically applied at high concentrations (≥ 10 μM) to silence ON pathways (Slaughter & Miller, 1981). The 50 nM dose delivered in our experiments likely bound only a small fraction of mGluR6 receptors, yet it effectively reduced pathological noise without blocking visual signals from healthy rods (fig. 6). RBCs are depolarized substantially following loss of glutamatergic input at just a fraction of rod synapses (fig. 8); a low dose of L-AP4 appears to reduce this pathological activity while still allowing RBCs to detect light-evoked changes in glutamate release at healthy synapses and transmit visual information to downstream targets. L-AP4 likely decreases the gain of healthy rod-RBC synapses, but by hyperpolarizing RBCs it also increases the gain of their synaptic outputs to AIIs (fig. 8). Consequently, signals originating in surviving rods are transmitted effectively through the circuit to RGCs atop reduced background noise (fig. 6). 50 nM L-AP4 reduced contrast responses in the youngest *rd10* RGCs, but no similar effect was seen in WTs. This result may be due to an unknown effect of manipulating glutamatergic input while rod degeneration is just beginning.

Effects of 50 nM L-AP4 were difficult to detect in RBCs (data not shown) but were clearly evident in ONα RGCs (fig. 6). Each ONα RGC collects signals deriving from ∼500 upstream RBCs (Dunn & Rieke, 2006) and thereby averages out variable signals to reveal the effects of 50 nM L-AP4.

Following photoreceptor death, mGluR6 (Gargini et al., 2007) and TRPM1channels (Gayet-Primo & Puthussery, 2015) eventually migrate within the plasma membrane towards the soma. Their functionality post-mislocalization remains unclear, a further argument that the optimal treatment window that best preserves retinal structural integrity is prior to total rod loss(Scalabrino et al., 2023).

### Therapeutic approaches to treat retinitis pigmentosa

Exciting progress has been made toward restoring light sensitivity to degenerated retinas, either by replenishing photoreceptors using stem cells (Mahato et al., 2020), or by conferring light sensitivity onto other neurons in the circuit (Kralik et al., 2022; Leemann & Kleinlogel, 2023; Rodgers et al., 2023; Sahel et al., 2021; van Wyk & Kleinlogel, 2023). Gene therapy effectively targets individual mutations (Bennett et al., 1996; Pang et al., 2008; Tun et al., 2023), but this approach faces the daunting challenge posed by hundreds of different mutations that can cause RP (S. Daiger et al., 2013; Hamel, 2006; McLaughlin et al., 1995; O’Neal & Luther, 2023).

Photoreceptor degeneration induces immediate morphological remodeling that alters retinal circuitry (Denlinger et al., 2020; Fariss et al., 2000; Jones et al., 2012; Jones & Marc, 2005; Karlen et al., 2020; Pfeiffer, Anderson, et al., 2020) and may compromise the efficacy of subsequent treatments. Clinical trial studies (Fenner et al., 2023; Sahel et al., 2021) and optogenetic approaches in mouse models of RP (Rodgers et al., 2023) suggest that vision may still be restored after total photoreceptor loss. Morphological remodeling, however, continues even after photoreceptor loss (Pfeiffer, Marc, et al., 2020), and would further impede therapies in later stages of the disease. Pharmacological therapy approaches that enhance circuit-wide inhibition or block gap junctions effectively reduce pathological activity (Biswas et al., 2014; Toychiev et al., 2013), but they interfere with other components of retinal circuitry that may cause off-target effects. Our approach suggests a broadly applicable treatment that reduces pathological noise without compromising surviving visual signals.

Unfortunately, we were unable to test the longitudinal effects of L-AP4 on morphological remodeling during RP because *rd10* retinas did not tolerate intraocular injections, even at injection volumes below 0.5 µL (fig. 7) (Hombrebueno et al., 2014; Miki et al., 2009). We suspect that the generally fragile state of the *rd10* retina largely contributed to its inability to withstand mild trauma caused by the injection. The small hole required to insert the syringe needle (30G) reduced the intraocular pressure, a change that was well tolerated by WT eyes but not by *rd10* eyes; species with larger vitreal volumes may better tolerate injections. Alternatively, less mechanically traumatic delivery methods, e.g., nanoparticles, ocular drug implants, or eyedrops (Batabyal et al., 2020; Meza-Rios et al., 2020; Sun et al., 2021), may preserve functional tissue.

## Acknowledgments

We thank Ms. Hua Tian for mouse husbandry, Dr. Jeff Bowen for helpful discussions regarding statistics, and Dr. Francisco Nadal-Nicolás for help with intraocular injections. This research was supported by the NINDS Intramural Research Program (NS003039 to JSD).

## References

Barhoum, R., Martínez-Navarrete, G., Corrochano, S., Germain, F., Fernandez-Sanchez, L., de la Rosa, E. J., de la Villa, P., & Cuenca, N. (2008). Functional and structural modifications during retinal degeneration in the rd10 mouse. Neuroscience, 155(3), 698–713. 10.1016/j.neuroscience.2008.06.042

Barone, I., Novelli, E., & Strettoi, E. (2014). Long-term preservation of cone photoreceptors and visual acuity in rd10 mutant mice exposed to continuous environmental enrichment. Molecular Vision, 20, 1545–1556.

Batabyal, S., Gajjeraman, S., Tchedre, K., Dibas, A., Wright, W., & Mohanty, S. (2020). Near-Infrared Laser-Based Spatially Targeted Nano-enhanced Optical Delivery of Therapeutic Genes to Degenerated Retina. Molecular Therapy. Methods & Clinical Development, 17, 758–770. 10.1016/j.omtm.2020.03.030

Bennett, J., Tanabe, T., Sun, D., Zeng, Y., Kjeldbye, H., Gouras, P., & Maguire, A. M. (1996). Photoreceptor cell rescue in retinal degeneration (rd) mice by in vivo gene therapy. Nature Medicine, 2(6), 649–654. 10.1038/nm0696-649

Biswas, S., Haselier, C., Mataruga, A., Thumann, G., Walter, P., & Müller, F. (2014). Pharmacological analysis of intrinsic neuronal oscillations in rd10 retina. PloS One, 9(6), e99075. 10.1371/journal.pone.0099075

Bleckert, A., Parker, E. D., Kang, Y., Pancaroglu, R., Soto, F., Lewis, R., Craig, A. M., & Wong, R. O. L. (2013). Spatial relationships between GABAergic and glutamatergic synapses on the dendrites of distinct types of mouse retinal ganglion cells across development. PloS One, 8(7), e69612. 10.1371/journal.pone.0069612

Bowes, C., Li, T., Danciger, M., Baxter, L. C., Applebury, M. L., & Farber, D. B. (1990). Retinal degeneration in the rd mouse is caused by a defect in the beta subunit of rod cGMP-phosphodiesterase. Nature, 347(6294), 677–680. 10.1038/347677a0

Busskamp, V., Duebel, J., Balya, D., Fradot, M., Viney, T. J., Siegert, S., Groner, A. C., Cabuy, E., Forster, V., Seeliger, M., Biel, M., Humphries, P., Paques, M., Mohand-Said, S., Trono, D., Deisseroth, K., Sahel, J. A., Picaud, S., & Roska, B. (2010). Genetic reactivation of cone photoreceptors restores visual responses in retinitis pigmentosa. Science (New York, N.Y.), 329(5990), 413–417. 10.1126/science.1190897

Chang, B., Hawes, N. L., Pardue, M. T., German, A. M., Hurd, R. E., Davisson, M. T., Nusinowitz, S., Rengarajan, K., Boyd, A. P., Sidney, S. S., Phillips, M. J., Stewart, R. E., Chaudhury, R., Nickerson, J. M., Heckenlively, J. R., & Boatright, J. H. (2007). Two mouse retinal degenerations caused by missense mutations in the beta-subunit of rod cGMP phosphodiesterase gene. Vision Research, 47(5), 624–633. 10.1016/j.visres.2006.11.020

Cross, N., van Steen, C., Zegaoui, Y., Satherley, A., & Angelillo, L. (2022). Current and Future Treatment of Retinitis Pigmentosa. Clinical Ophthalmology (Auckland, N.Z.), 16, 2909– 2921. 10.2147/OPTH.S370032

Daiger, S. P., Bowne, S. J., & Sullivan, L. S. (2007). Perspective on Genes and Mutations Causing Retinitis Pigmentosa. Archives of Ophthalmology, 125(2), 151–158. 10.1001/archopht.125.2.151

Daiger, S., Sullivan, L., & Bowne, S. (2013). Genes and mutations causing retinitis pigmentosa. Clinical Genetics, 84(2), 10.1111/cge.12203. https://doi.org/10.1111/cge.12203

Denlinger, B., Helft, Z., Telias, M., Lorach, H., Palanker, D., & Kramer, R. H. (2020). Local photoreceptor degeneration causes local pathophysiological remodeling of retinal neurons. JCI Insight, 5(2), e132114, 132114. 10.1172/jci.insight.132114

Dunn, F. A., & Rieke, F. (2006). The impact of photoreceptor noise on retinal gain controls. Current Opinion in Neurobiology, 16(4), 363–370. 10.1016/j.conb.2006.06.013

Fariss, R. N., Li, Z. Y., & Milam, A. H. (2000). Abnormalities in rod photoreceptors, amacrine cells, and horizontal cells in human retinas with retinitis pigmentosa. American Journal of Ophthalmology, 129(2), 215–223. 10.1016/s0002-9394(99)00401-8

Fenner, B. J., Jamshidi, F., Bhuyan, R., Fortenbach, C. R., Jin, H. D., Boyce, T. M., Binkley, E. M., Han, I. C., Sohn, E. H., Boldt, H. C., Folk, J. C., Russell, S. R., Stone, E. M., & Russell, J. F. (2023). Vitreoretinal procedures in patients with inherited retinal disease. Ophthalmology. Retina, S2468–6530(23)00571-7. 10.1016/j.oret.2023.10.020

Franke, K., Maia Chagas, A., Zhao, Z., Zimmermann, M. J., Bartel, P., Qiu, Y., Szatko, K. P., Baden, T., & Euler, T. (2019). An arbitrary-spectrum spatial visual stimulator for vision research. eLife, 8, e48779. 10.7554/eLife.48779

Gargini, C., Terzibasi, E., Mazzoni, F., & Strettoi, E. (2007). IRetinal Organization in the retinal degeneration 10 (rd10) Mutant Mouse: A Morphological and ERG Study. The Journal of Comparative Neurology, 500(2), 222–238. 10.1002/cne.21144

Gayet-Primo, J., & Puthussery, T. (2015). Alterations in Kainate Receptor and TRPM1 Localization in Bipolar Cells after Retinal Photoreceptor Degeneration. Frontiers in Cellular Neuroscience, 9. 10.3389/fncel.2015.00486

Goetz, J., Jessen, Z. F., Jacobi, A., Mani, A., Cooler, S., Greer, D., Kadri, S., Segal, J., Shekhar, K., Sanes, J. R., & Schwartz, G. W. (2022). Unified classification of mouse retinal ganglion cells using function, morphology, and gene expression. Cell Reports, 40(2), 111040. 10.1016/j.celrep.2022.111040

Goo, Y. S., Park, D. J., Ahn, J. R., & Senok, S. S. (2015). Spontaneous Oscillatory Rhythms in the Degenerating Mouse Retina Modulate Retinal Ganglion Cell Responses to Electrical Stimulation. Frontiers in Cellular Neuroscience, 9, 512. 10.3389/fncel.2015.00512

Govardovskii, V. I., Fyhrquist, N., Reuter, T., Kuzmin, D. G., & Donner, K. (2000). In search of the visual pigment template. Visual Neuroscience, 17(4), 509–528. 10.1017/s0952523800174036

Graydon, C. W., Lieberman, E. E., Rho, N., Briggman, K. L., Singer, J. H., & Diamond, J. S. (2018). Synaptic Transfer between Rod and Cone Pathways Mediated by AII Amacrine Cells in the Mouse Retina. Current Biology: CB, 28(17), 2739–2751.e3. 10.1016/j.cub.2018.06.063

Grimes, W. N., Schwartz, G. W., & Rieke, F. (2014). The synaptic and circuit mechanisms underlying a change in spatial encoding in the retina. Neuron, 82(2), 460–473. 10.1016/j.neuron.2014.02.037

Hamel, C. (2006). Retinitis pigmentosa. Orphanet Journal of Rare Diseases, 1, 40. 10.1186/1750-1172-1-40

Herrmann, R., Heflin, S. J., Hammond, T., Lee, B., Wang, J., Gainetdinov, R. R., Caron, M. G., Eggers, E. D., Frishman, L. J., McCall, M. A., & Arshavsky, V. Y. (2011). Rod Vision Is Controlled by Dopamine-Dependent Sensitization of Rod Bipolar Cells by GABA. Neuron, 72(1), 101–110. 10.1016/j.neuron.2011.07.030

Hombrebueno, J. R., Luo, C., Guo, L., Chen, M., & Xu, H. (2014). Intravitreal Injection of Normal Saline Induces Retinal Degeneration in the C57BL/6J Mouse. Translational Vision Science & Technology, 3(2), 3. 10.1167/tvst.3.2.3

Jae, S. A., Ahn, K. N., Kim, J. Y., Seo, J. H., Kim, H. K., & Goo, Y. S. (2013). Electrophysiological and Histologic Evaluation of the Time Course of Retinal Degeneration in the rd10 Mouse Model of Retinitis Pigmentosa. The Korean Journal of Physiology & Pharmacology: Official Journal of the Korean Physiological Society and the Korean Society of Pharmacology, 17(3), 229–235. 10.4196/kjpp.2013.17.3.229

Jones, B. W., Kondo, M., Terasaki, H., Lin, Y., McCall, M., & Marc, R. E. (2012). Retinal remodeling. Japanese Journal of Ophthalmology, 56(4), 289–306. 10.1007/s10384-012-0147-2

Jones, B. W., & Marc, R. E. (2005). Retinal remodeling during retinal degeneration. Experimental Eye Research, 81(2), 123–137. 10.1016/j.exer.2005.03.006

Karlen, S. J., Miller, E. B., & Burns, M. E. (2020). Microglia Activation and Inflammation During the Death of Mammalian Photoreceptors. Annual Review of Vision Science, 6, 149–169. 10.1146/annurev-vision-121219-081730

Kim, T.-H., Son, T., Lu, Y., Alam, M., & Yao, X. (2018). Comparative Optical Coherence Tomography Angiography of Wild-Type and rd10 Mouse Retinas. Translational Vision Science & Technology, 7(6), 42. 10.1167/tvst.7.6.42

Kim, Y. N., Song, J. S., Oh, S. H., Kim, Y. J., Yoon, Y. H., Seo, E.-J., Seol, C. A., Lee, S.-M., Choi, J.-M., Seo, G. H., Keum, C., Lee, B. H., & Lee, J. Y. (2020). Clinical characteristics and disease progression of retinitis pigmentosa associated with PDE6B mutations in Korean patients. Scientific Reports, 10(1), Article 1. 10.1038/s41598-020-75902-z

Kralik, J., van Wyk, M., Stocker, N., & Kleinlogel, S. (2022). Bipolar cell targeted optogenetic gene therapy restores parallel retinal signaling and high-level vision in the degenerated retina. Communications Biology, 5(1), Article 1. 10.1038/s42003-022-04016-1

Krieger, B., Qiao, M., Rousso, D. L., Sanes, J. R., & Meister, M. (2017). Four alpha ganglion cell types in mouse retina: Function, structure, and molecular signatures. PloS One, 12(7), e0180091. 10.1371/journal.pone.0180091

Kuehlewein, L., Zobor, D., Stingl, K., Kempf, M., Nasser, F., Bernd, A., Biskup, S., Cremers, F. P. M., Khan, M. I., Mazzola, P., Schäferhoff, K., Heinrich, T., Haack, T. B., Wissinger, B., Zrenner, E., Weisschuh, N., & Kohl, S. (2021). Clinical Phenotype of PDE6B-Associated Retinitis Pigmentosa. International Journal of Molecular Sciences, 22(5), 2374. 10.3390/ijms22052374

Laboissonniere, L. A., Goetz, J. J., Martin, G. M., Bi, R., Lund, T. J. S., Ellson, L., Lynch, M. R., Mooney, B., Wickham, H., Liu, P., Schwartz, G. W., & Trimarchi, J. M. (2019). Molecular signatures of retinal ganglion cells revealed through single cell profiling. Scientific Reports, 9(1), 15778. 10.1038/s41598-019-52215-4

Lee, J. Y., Care, R. A., Della Santina, L., & Dunn, F. A. (2021). Impact of Photoreceptor Loss on Retinal Circuitry. Annual Review of Vision Science, 7, 105–128. 10.1146/annurev-vision-100119-124713

Leemann, S., & Kleinlogel, S. (2023). Functional optimization of light-activatable Opto-GPCRs: Illuminating the importance of the proximal C-terminus in G-protein specificity. Frontiers in Cell and Developmental Biology, 11, 1053022. 10.3389/fcell.2023.1053022

Li, Y., Cohen, E. D., & Qian, H. (2020). Rod and Cone Coupling Modulates Photopic ERG Responses in the Mouse Retina. Frontiers in Cellular Neuroscience, 14, 566712. 10.3389/fncel.2020.566712

Li, Y., Zhang, Y., Chen, S., Vernon, G., Wong, W. T., & Qian, H. (2018). Light-Dependent OCT Structure Changes in Photoreceptor Degenerative rd 10 Mouse Retina. Investigative Ophthalmology & Visual Science, 59(2), 1084–1094. 10.1167/iovs.17-23011

Lyubarsky, A. L., Daniele, L. L., & Pugh, E. N. (2004). From candelas to photoisomerizations in the mouse eye by rhodopsin bleaching in situ and the light-rearing dependence of the major components of the mouse ERG. Vision Research, 44(28), 3235–3251. 10.1016/j.visres.2004.09.019

Mahato, B., Kaya, K. D., Fan, Y., Sumien, N., Shetty, R. A., Zhang, W., Davis, D., Mock, T., Batabyal, S., Ni, A., Mohanty, S., Han, Z., Farjo, R., Forster, M. J., Swaroop, A., & Chavala, S. H. (2020). Pharmacologic fibroblast reprogramming into photoreceptors restores vision. Nature, 581(7806), Article 7806. 10.1038/s41586-020-2201-4

McLaughlin, M. E., Ehrhart, T. L., Berson, E. L., & Dryja, T. P. (1995). Mutation spectrum of the gene encoding the beta subunit of rod phosphodiesterase among patients with autosomal recessive retinitis pigmentosa. Proceedings of the National Academy of Sciences of the United States of America, 92(8), 3249–3253. 10.1073/pnas.92.8.3249

Meza-Rios, A., Navarro-Partida, J., Armendariz-Borunda, J., & Santos, A. (2020). Therapies Based on Nanoparticles for Eye Drug Delivery. Ophthalmology and Therapy. 10.1007/s40123-020-00257-7

Miki, K., Miki, A., Matsuoka, M., Muramatsu, D., Hackett, S. F., & Campochiaro, P. A. (2009). Effects of Intraocular Ranibizumab and Bevacizumab in Transgenic Mice Expressing Human Vascular Endothelial Growth Factor. Ophthalmology, 116(9), 1748–1754. 10.1016/j.ophtha.2009.05.020

Naples, M. A., & Hampson, D. R. (2001). Pharmacological profiles of the metabotropic glutamate receptor ligands [3H]L-AP4 and [3H]CPPG. Neuropharmacology, 40(2), 170–177. 10.1016/S0028-3908(00)00128-3

Narayan, D. S., Wood, J. P. M., Chidlow, G., & Casson, R. J. (2016). A review of the mechanisms of cone degeneration in retinitis pigmentosa. Acta Ophthalmologica, 94(8), 748–754. 10.1111/aos.13141

Nath, A., Grimes, W. N., & Diamond, J. S. (2023). Layers of inhibitory networks shape receptive field properties of AII amacrine cells. Cell Reports, 42(11), 113390. 10.1016/j.celrep.2023.113390

Neher, E. (2015). Merits and Limitations of Vesicle Pool Models in View of Heterogeneous Populations of Synaptic Vesicles. Neuron, 87(6), 1131–1142. 10.1016/j.neuron.2015.08.038

Oesch, N. W., & Diamond, J. S. (2011). Ribbon synapses compute temporal contrast and encode luminance in retinal rod bipolar cells. Nature Neuroscience, 14(12), Article 12. 10.1038/nn.2945

Oltedal, L., Veruki, M. L., & Hartveit, E. (2009). Passive membrane properties and electrotonic signal processing in retinal rod bipolar cells. The Journal of Physiology, 587(Pt 4), 829–849. 10.1113/jphysiol.2008.165415

O’Neal, T. B., & Luther, E. E. (2023). Retinitis Pigmentosa. In StatPearls. StatPearls Publishing. http://www.ncbi.nlm.nih.gov/books/NBK519518/

Pang, J.-J., Boye, S. L., Kumar, A., Dinculescu, A., Deng, W., Li, J., Li, Q., Rani, A., Foster, T. C., Chang, B., Hawes, N. L., Boatright, J. H., & Hauswirth, W. W. (2008). AAV-mediated gene therapy for retinal degeneration in the rd10 mouse containing a recessive PDEbeta mutation. Investigative Ophthalmology & Visual Science, 49(10), 4278–4283. 10.1167/iovs.07-1622

Pennesi, M. E., Michaels, K. V., Magee, S. S., Maricle, A., Davin, S. P., Garg, A. K., Gale, M. J., Tu, D. C., Wen, Y., Erker, L. R., & Francis, P. J. (2012). Long-term characterization of retinal degeneration in rd1 and rd10 mice using spectral domain optical coherence tomography. Investigative Ophthalmology & Visual Science, 53(8), 4644–4656. 10.1167/iovs.12-9611

Perlman, I. (1995). The Electroretinogram: ERG. In H. Kolb, E. Fernandez, & R. Nelson (Eds.), Webvision: The Organization of the Retina and Visual System. University of Utah Health Sciences Center. http://www.ncbi.nlm.nih.gov/books/NBK11554/

Pfeiffer, R. L., Anderson, J. R., Dahal, J., Garcia, J. C., Yang, J.-H., Sigulinsky, C. L., Rapp, K., Emrich, D. P., Watt, C. B., Johnstun, H. A., Houser, A. R., Marc, R. E., & Jones, B. W. (2020). A pathoconnectome of early neurodegeneration: Network changes in retinal degeneration. Experimental Eye Research, 199, 108196. 10.1016/j.exer.2020.108196

Pfeiffer, R. L., Marc, R. E., & Jones, B. W. (2020). Persistent remodeling and neurodegeneration in late-stage retinal degeneration. Progress in Retinal and Eye Research, 74, 100771. 10.1016/j.preteyeres.2019.07.004

Phillips, M. J., Otteson, D. C., & Sherry, D. M. (2010). Progression of neuronal and synaptic remodeling in the rd10 mouse model of retinitis pigmentosa. The Journal of Comparative Neurology, 518(11), 2071–2089. 10.1002/cne.22322

Pinto, L. H., Invergo, B., Shimomura, K., Takahashi, J. S., & Troy, J. B. (2007). Interpretation of the mouse electroretinogram. Documenta Ophthalmologica, 115(3), 127–136. 10.1007/s10633-007-9064-y

Punzo, C., Kornacker, K., & Cepko, C. L. (2009). Stimulation of the insulin/mTOR pathway delays cone death in a mouse model of retinitis pigmentosa. Nature Neuroscience, 12(1), 44–52. 10.1038/nn.2234

Puthussery, T., Gayet-Primo, J., Pandey, S., Duvoisin, R. M., & Taylor, W. R. (2009). Differential loss and preservation of glutamate receptor function in bipolar cells in the rd10 mouse model of retinitis pigmentosa. The European Journal of Neuroscience, 29(8), 1533–1542. 10.1111/j.1460-9568.2009.06728.x

Puthussery, T., & Taylor, W. R. (2010). Functional Changes in Inner Retinal Neurons in Animal Models of Photoreceptor Degeneration. In R. E. Anderson, J. G. Hollyfield, & M. M. LaVail (Eds.), Retinal Degenerative Diseases: Laboratory and Therapeutic Investigations (pp. 525–532). Springer. 10.1007/978-1-4419-1399-9_60

Rodgers, J., Hughes, S., Lindner, M., Allen, A. E., Ebrahimi, A. S., Storchi, R., Peirson, S. N., Lucas, R. J., & Hankins, M. W. (2023). Functional integrity of visual coding following advanced photoreceptor degeneration. Current Biology, 33(3), 474–486.e5. 10.1016/j.cub.2022.12.026

Sahel, J.-A., Boulanger-Scemama, E., Pagot, C., Arleo, A., Galluppi, F., Martel, J. N., Esposti, S. D., Delaux, A., de Saint Aubert, J.-B., de Montleau, C., Gutman, E., Audo, I., Duebel, J., Picaud, S., Dalkara, D., Blouin, L., Taiel, M., & Roska, B. (2021). Partial recovery of visual function in a blind patient after optogenetic therapy. Nature Medicine, 27(7), Article 7. 10.1038/s41591-021-01351-4

Scalabrino, M. L., Thapa, M., Wang, T., Sampath, A. P., Chen, J., & Field, G. D. (2023). Late gene therapy limits the restoration of retinal function in a mouse model of retinitis pigmentosa. Nature Communications, 14(1), 8256. 10.1038/s41467-023-44063-8

Schneeweis, D. M., & Schnapf, J. L. (1995). Photovoltage of rods and cones in the macaque retina. Science (New York, N.Y.), 268(5213), 1053–1056. 10.1126/science.7754386

Schwartz, G. W., Okawa, H., Dunn, F. A., Morgan, J. L., Kerschensteiner, D., Wong, R. O., & Rieke, F. (2012). The spatial structure of a nonlinear receptive field. Nature Neuroscience, 15(11), Article 11. 10.1038/nn.3225

Simon, C.-J., Sahel, J.-A., Duebel, J., Herlitze, S., & Dalkara, D. (2020). Opsins for vision restoration. Biochemical and Biophysical Research Communications, 527(2), 325–330. 10.1016/j.bbrc.2019.12.117

Singer, J. H., & Diamond, J. S. (2003). Sustained Ca2+ entry elicits transient postsynaptic currents at a retinal ribbon synapse. The Journal of Neuroscience: The Official Journal of the Society for Neuroscience, 23(34), 10923–10933. 10.1523/JNEUROSCI.23-34-10923.2003

Singer, J. H., & Diamond, J. S. (2006). Vesicle depletion and synaptic depression at a mammalian ribbon synapse. Journal of Neurophysiology, 95(5), 3191–3198. 10.1152/jn.01309.2005

Slaughter, M. M., & Miller, R. F. (1981). 2-amino-4-phosphonobutyric acid: A new pharmacological tool for retina research. Science (New York, N.Y.), 211(4478), 182–185. 10.1126/science.6255566

Stasheff, S. F., Shankar, M., & Andrews, M. P. (2011). Developmental time course distinguishes changes in spontaneous and light-evoked retinal ganglion cell activity in rd1 and rd10 mice. Journal of Neurophysiology, 105(6), 3002–3009. 10.1152/jn.00704.2010

Stefanov, A., Novelli, E., & Strettoi, E. (2020). Inner retinal preservation in the photoinducible I307N rhodopsin mutant mouse, a model of autosomal dominant retinitis pigmentosa. Journal of Comparative Neurology, 528(9), 1502–1522. 10.1002/cne.24838

Sun, Y. J., Lin, C.-H., Wu, M.-R., Lee, S. H., Yang, J., Kunchur, C. R., Mujica, E. M., Chiang, B., Jung, Y. S., Wang, S., & Mahajan, V. B. (2021). An intravitreal implant injection method for sustained drug delivery into mouse eyes. Cell Reports Methods, 1(8), 100125. 10.1016/j.crmeth.2021.100125

Thomsen, C. (1997). The l-AP4 receptor. General Pharmacology: The Vascular System, 29(2), 151–158. 10.1016/S0306-3623(96)00417-X

Toychiev, A. H., Ivanova, E., Yee, C. W., & Sagdullaev, B. T. (2013). Block of Gap Junctions Eliminates Aberrant Activity and Restores Light Responses during Retinal Degeneration. Journal of Neuroscience, 33(35), 13972–13977. 10.1523/JNEUROSCI.2399-13.2013

Trenholm, S., & Awatramani, G. B. (2015). Origins of spontaneous activity in the degenerating retina. Frontiers in Cellular Neuroscience, 9. https://www.frontiersin.org/articles/10.3389/fncel.2015.00277

Tun, S. B. B., Shepherdson, E., Tay, H. G., & Barathi, V. A. (2023). Sub-Retinal Delivery of Human Embryonic Stem Cell Derived Photoreceptor Progenitors in rd10 Mice. Journal of Visualized Experiments: JoVE, 200. 10.3791/65848

van Wyk, M., & Kleinlogel, S. (2023). A visual opsin from jellyfish enables precise temporal control of G protein signalling. Nature Communications, 14(1), 2450. 10.1038/s41467-023-38231-z

Völgyi, B., Deans, M. R., Paul, D. L., & Bloomfield, S. A. (2004). Convergence and Segregation of the Multiple Rod Pathways in Mammalian Retina. The Journal of Neuroscience, 24(49), 11182–11192. 10.1523/JNEUROSCI.3096-04.2004

Wang, B.-Y., Chen, Z. C., Bhuckory, M., Huang, T., Shin, A., Zuckerman, V., Ho, E., Rosenfeld, E., Galambos, L., Kamins, T., Mathieson, K., & Palanker, D. (2022). Electronic photoreceptors enable prosthetic visual acuity matching the natural resolution in rats. Nature Communications, 13(1), 6627. 10.1038/s41467-022-34353-y

Wang, T., Pahlberg, J., Cafaro, J., Frederiksen, R., Cooper, A. J., Sampath, A. P., Field, G. D., & Chen, J. (2019). Activation of Rod Input in a Model of Retinal Degeneration Reverses Retinal Remodeling and Induces Formation of Functional Synapses and Recovery of Visual Signaling in the Adult Retina. Journal of Neuroscience, 39(34), 6798–6810. 10.1523/JNEUROSCI.2902-18.2019

Yu, C., Roubeix, C., Sennlaub, F., & Saban, D. R. (2020). Microglia versus Monocytes: Distinct Roles in Degenerative Diseases of the Retina. Trends in Neurosciences, 43(6), 433–449. 10.1016/j.tins.2020.03.012

